# Mitochondrial fission process 1 (MTFP1) controls bioenergetic efficiency and prevents inflammatory cardiomyopathy and heart failure in mice

**DOI:** 10.1101/2021.10.21.465262

**Authors:** Erminia Donnarumma, Michael Kohlhaas, Elodie Vimont, Etienne Kornobis, Thibault Chaze, Quentin Giai Gianetto, Mariette Matondo, Maryse Moya-Nilges, Christoph Maack, Timothy Wai

## Abstract

Mitochondria are paramount to the metabolism and survival of cardiomyocytes. Here we show that Mitochondrial Fission Process 1 (MTFP1) is essential for cardiac structure and function. Constitutive knockout of cardiomyocyte MTFP1 in mice resulted in adult-onset dilated cardiomyopathy (DCM) characterized by sterile inflammation and cardiac fibrosis that progressed to heart failure and middle-aged death. Failing hearts from cardiomyocyte-restricted knockout mice displayed a general decline in mitochondrial gene expression and oxidative phosphorylation (OXPHOS) activity. Pre-DCM, we observed no defects in mitochondrial morphology, content, gene expression, OXPHOS assembly nor phosphorylation dependent respiration. However, knockout cardiac mitochondria displayed reduced membrane potential and increased non-phosphorylation dependent respiration, which could be rescued by pharmacological inhibition of the adenine nucleotide translocase ANT. Primary cardiomyocytes from pre-symptomatic knockout mice exhibited normal excitation-contraction coupling but increased sensitivity to programmed cell death (PCD), which was accompanied by an opening of the mitochondrial permeability transition pore (mPTP). Intriguingly, mouse embryonic fibroblasts deleted for *Mtfp1* recapitulated PCD sensitivity and mPTP opening, both of which could be rescued by pharmacological or genetic inhibition of the mPTP regulator Cyclophilin D. Collectively, our data demonstrate that contrary to previous in vitro studies, the loss of the MTFP1 promotes mitochondrial uncoupling and increases cell death sensitivity, causally mediating pathogenic cardiac remodeling.

## Introduction

Mitochondria are multifaceted organelles that are essential in every tissue of the body and are most abundant in the heart, where they control the metabolism and survival of cardiac cells (Kolwicz et al., 2013). Mitochondria are double membrane-bound organelles, composed of an inner (IMM) and an outer mitochondrial membrane (OMM), which separate the intermembrane space (IMS) from the matrix. The IMM extends internally to form cristae, which harbor essential macromolecular complexes such as the machinery of oxidative phosphorylation (OXPHOS). The OXPHOS system is comprised of two functional entities: the electron transport chain (ETC) and the phosphorylation system, which includes the integral membrane ATP synthase and carriers such as the adenine nucleotide translocase (ANT), which imports ADP into the matrix and exports ATP (Chance and Williams, 1956). The ETC is composed of four macromolecular complexes (I, II, III, and IV) and of mobile electrons carriers as coenzyme Q (CoQ) and cytochrome-c (Cyt *c*). The energy available for ATP synthesis is directly derived from the membrane potential (ΔΨ) and proton motive force generated across the IMM by electron transfer from by the ETC, which is then harnessed by the ATP synthase (complex V) to generate ATP (Mitchell, 1961). Continuous generation of ATP is fundamental for the function of cardiomyocytes, enabling them to meet the enormous energy requirement for the contraction–relaxation cycles that drive their contractility (Kohlhaas et al., 2017). Defects that impair OXPHOS assembly and function can promote fatal cardiomyopathies (Antonicka et al., 2003a, 2003b; Graham et al., 1997; Hansson et al., 2004; Karamanlidis et al., 2013). The beating heart demands near-maximum OXPHOS capacity, with scant aerobic ATP reserves under normal conditions (Mootha et al., 1997). Consequently, even modest uncoupling of the ETC from ATP synthesis, which can occur when protons are diverted back across the IMM through uncoupling channels rather than being used by the ATP synthase, would be expected to yield fatal consequences for cardiac function and health, although this has never been tested.

While most famous for their role as the powerhouse of the cell, mitochondria have proven to be essential for cardiac homeostasis through the regulation of various biosynthetic and signaling functions beyond OXPHOS, such as calcium buffering, reactive oxygen species (ROS) generation and maintenance, programmed cell death (PCD) and innate immune responses (Zhou and Tian, 2018). Under stress conditions, extrinsic or intrinsic signals can lead to a permeabilization of the outer membrane (MOMP) resulting in activation of caspases and PCD (Bock and Tait, 2020). At the IMM, long-lasting opening of the mitochondrial permeability transition pore (mPTP) allows for unselective diffusion of low molecular weight solutes and water (<1.5 kDa), causing an osmotic pressure in the matrix that causes mitochondrial swelling (Carraro et al., 2020) and rupture, thereby releasing pro-apoptotic and pro-inflammatory mitochondrial factors into the cytosol (Bock and Tait, 2020). While the molecular identity of the mPTP is still the subject of feverous debate, to date several factors have been identified to be unequivocally crucial for its activation. Cyclophilin D (CypD, encoded by *Ppif*), a mitochondrial matrix isomerase which can be inhibited pharmacologically by cyclosporin A (CsA), calcium overload and ROS are known to promote mPTP opening and PCD induction (Baines et al., 2005; Carraro et al., 2020; Nakagawa et al., 2005). Notably, ablation of CypD or treatment with CsA can protect animals from cardiomyocyte death and cardiomyopathy induced by genetic (Song et al., 2015), infectious (Milduberger et al., 2021), and surgical lesions (Nakagawa et al., 2005).

The maintenance of mitochondrial morphology and structure is of critical importance for cardiac function (Sprenger and Langer, 2019). Mitochondrial morphology is regulated by opposing forces of mitochondrial fusion and division, which must be tightly regulated to ensure organellar function and quality control (Giacomello et al., 2020; Ng et al., 2021). Intrinsic (Ashrafian et al., 2010; Chen et al., 2015; Kageyama et al., 2014; Song et al., 2015; Wai et al., 2015) or extrinsic (Acin-Perez et al., 2018; Chen et al., 2009; Shirakabe et al., 2016) lesions that upset the balance between mitochondrial fission and fusion have devastating consequences for cardiomyocyte function and cardiac health. Mitochondrial dynamics is orchestrated by dynamin-like GTPases: OPA1 and mitofusins (MFN1/2) execute inner and outer membrane fusion, respectively, while DRP1 performs mitochondrial constriction and division once recruited to the OMM via interactions that require integral membrane proteins and receptors, such as MFF, MiD49/51, and FIS1. DRP1 is the lynchpin of the mitochondrial fission apparatus, which is triggered to coalesce at contact sites with the ER, lysosomes, trans-golgi network, and actin by signals that can originate both outside (Cretin et al., 2021; Giacomello et al., 2020) and inside (Cho et al., 2017; Lewis et al., 2016; Tondera, 2005) mitochondria.

While it is unclear how IMM fission is executed, the aptly-named inner membrane protein Mitochondrial fission process 1 (MTFP1/MTP18), has emerged as a promising scaffold for the IMM division apparatus, whose formal identification remains elusive (Ng et al., 2021). As per its namesake, *Mtfp1* was initially identified as a gene whose ablation was reported to reduce mitochondrial fission, as well as the proliferation and viability of cultured cells (Tondera, 2005; Tondera et al., 2004). The pro-apoptotic effects of *Mtfp1* depletion have been replicated in various cell lines (Duroux-Richard et al., 2016; Wang et al., 2017), however, more recent studies by the Li group (Aung et al., 2017a, 2017b; Wang et al., 2017) have reported the contrary in cultured cells, thus fueling the existing narrative that elongated mitochondria resulting from MTFP1 depletion protects cells from PCD. Whether this paradigm holds true in vivo has never been explored.

In this study, we created a cardiomyocyte-specific *Mtfp1* knockout mouse model to specifically investigate the role of this protein in vivo. Contrary to previous in vitro studies (Wang et al., 2017), we show that MTFP1 plays an essential role in maintaining cardiac energy metabolism as its deletion in post-natal cardiomyocytes drives a progressive dilated cardiomyopathy (DCM) culminating in HF and middle-aged death in mice. Surprisingly, MTFP1 ablation does not appreciably alter mitochondrial morphology in the heart and is entirely dispensable for mitochondrial fission. Unexpectedly, we discovered that MTFP1 depletion reduces OXPHOS efficiency in cardiac mitochondria by increasing proton leak through the adenine nucleotide translocase (ANT). Finally, we show MTFP1 ablation increases mPTP opening and renders cardiomyocytes and embryonic fibroblasts more sensitive to PCD. Altogether, our data reveal an unexpected role of MTFP1 in mitochondrial bioenergetics and provide mechanistic insights into how MTFP1 regulates the life and death of the cell.

## Results

### *Mtfp1* deletion in cardiomyocytes causes dilated cardiomyopathy and middle-aged death

MTFP1 is predicted to localize in mitochondrial inner membrane (Figure 1A), and *MTFP1* is highly expressed in human cardiac tissue (GTEx plot, Figure S1A). To investigate the importance of MTFP1 for cardiac function we began by confirming its expression and submitochondrial localization in the mouse heart. Protease protection assays and alkaline carbonate extraction experiments performed on isolated cardiac mitochondria allowed us to demonstrate the inner membrane localization of MTFP1 in cardiac mitochondria (Figure S1B-C). To investigate its role in vivo, we generated a conditional mouse model for *Mtfp1* (*Mtfp1^LoxP/LoxP^*) on a C57Bl6/N background. Conditional mice were crossed with transgenic mice expressing Cre recombinase under the control of alpha myosin heavy chain promoter (Agah et al., 1997) (Myh6) to specifically ablate MTFP1 in post-natal cardiomyocytes (Figure 1B, S1D). We previously showed that genetic deletion mediated by Myh6-Cre occurs during the perinatal period (Wai et al., 2015) and by 8 weeks of age cardiomyocyte-specific *Mtfp1* KO mice (Myh6-Cre^Tg/+^ *Mtfp1 ^LoxP/LoxP^*, cMKO mice) exhibited a 7-fold reduction in *Mtfp1* mRNA (Figure 1C) and undetectable levels of MTFP1 protein in cardiac lysates assessed by immunoblot analysis (Figure 1D) and shotgun proteomics (Figure 1E, Dataset EV1). Both male and female cMKO mice were generated according to Mendelian proportions and were outwardly normal and viable. However, cMKO mice had significantly shortened lifespans [median life span: 26.4 weeks (male), 37.5 weeks (female)] relative to wild type (WT) littermates (Figure 1F, Figure S1E), demonstrating that MTFP1 is required to protect against middle-aged death.

**Figure 1.**
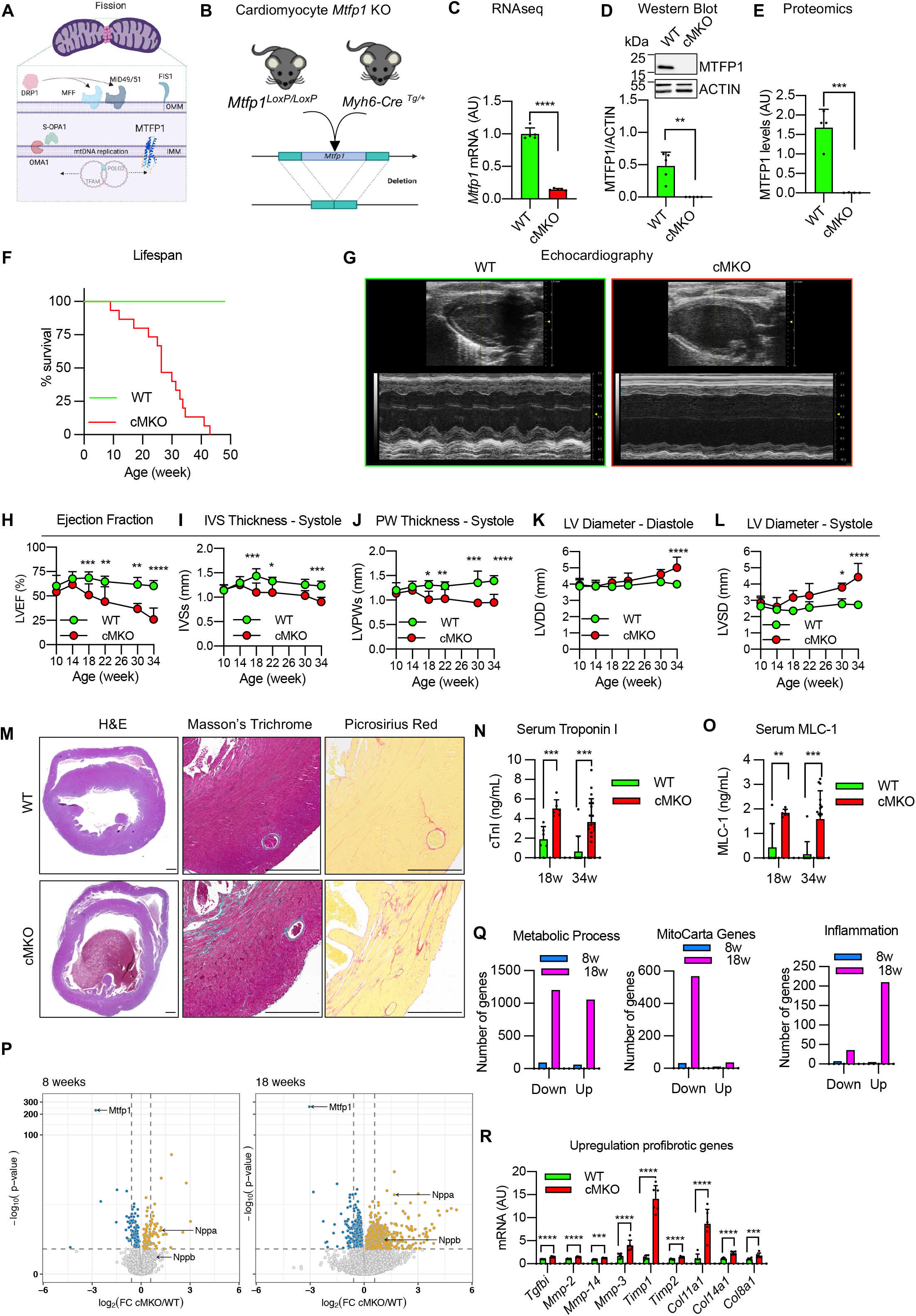
*Mtfp1* deletion in cardiomyocytes causes dilated cardiomyopathy and middle-aged death in mice. **A)** AlphaFold prediction of MTFP1 as integral protein of the inner mitochondrial membrane (IMM) associated with mitochondrial fission. DRP1 binds to its receptors MFF or MiD49/51 to initiate mitochondrial constriction. IMM fission occurs coincidently with mtDNA replication mediated by TFAM and POLG2. S-OPA1 accumulation generated by OMA1 accelerates fission. **B)** Generation of a cardiomyocyte-specific *Mtfp1* KO (cMKO) mouse model. **C)** *Mtfp1* mRNA expression measured by bulk RNAseq (arbitrary units; AU) in heart tissue of WT (n=5) and cMKO (n=5) mice at 8 weeks (Dataset EV2). Data represent mean ± SD; 2-tailed unpaired Student’s t-test, ****p<0.0001. **D)** Densitometric quantification of immunoblot analysis of cardiac lysates from WT (n=5) and cMKO (n=5) mice at 8 weeks using the indicated antibodies. Data represent mean ± SD; Unpaired t-test, **p<0.01. **E)** MTFP1 protein expression in cardiac tissue of WT (n=4) and cMKO (n=4) at 18 weeks measured by label free quantification (LFQ) mass spectrometry (MS) (Dataset EV1). Data represent mean ± SD; 2-tailed unpaired Student’s t-test, ***p<0.001. **F)** Kaplan-Meier survival curve of WT (n=9) and cMKO (n=15) male mice. Median lifespan of cMKO mice is 26.4 weeks. Log-rank (Mantel-Cox) test, ****p < 0.0001. **G-L)** Longitudinal echocardiography of WT and cMKO mice from 10 to 34 weeks of age. **G)** Representative M-Mode echocardiographic images of left ventricles from WT (left) and cMKO (right) of male mice at 34 weeks. **H)** Quantification of left ventricular ejection fraction (% LVEF) of WT and cMKO mice at indicated ages. Data represent mean ± SD; 2way Anova - Sidak’s multiple comparison test: 10 week WT (n=13) vs cMKO (n=18); 14 week WT (n=4) vs cMKO (n=6); 18 week WT (n=10) vs cMKO (n=13) ***p<0.001; 22 week WT (n=4) vs cMKO (n=6) **p<0.01; 30 week WT (n=3) vs cMKO (n=7) **p<0.01; 34 week WT (n=4) vs cMKO (n=10)****p<0.0001. **I)** Quantification of systolic interventricular septum thickness (IVSs, mm) of WT and cMKO mice at indicated ages. Data represent mean ± SD; 2way Anova - Sidak’s multiple comparison test, 18 weeks ***p<0.001; 22 weeks *p<0.05; 34 weeks ***p<0.001. **J)** Quantification of left ventricle posterior wall thickness at systole (LVPWs, mm) of WT and cMKO mice at indicated ages. Data represent mean ± SD; 2way Anova - Sidak’s multiple comparison test, 18 weeks **p<0.01; 22 weeks *p<0.05; 30 weeks ***p<0.001, 34 weeks ****p<0.0001. **K)** Quantification of left ventricle end diastolic diameter (LVDD, mm) of WT and cMKO mice at indicated ages. Data represent mean ± SD; 2way Anova - Sidak’s multiple comparison test, 34 week ****p<0.0001. **L)** Quantification of left ventricle end systolic diameter (LVSD, mm) of WT and cMKO mice at indicated ages. Data represent mean ± SD; 2way Anova - Sidak’s multiple comparison test, 30 weeks *p<0.05; 34 weeks ***p<0.001. **M)** Representative histological images of cardiac short axis view of WT (n=4) and cMKO (n=4) at 34 weeks. H&E (left), Massons’s Trichrome (middle) and Picrosirius red (right) staining show cardiac remodeling and collagen deposition within the myocardium of cMKO mice. Scale bar 500 µM. **N)** Circulating levels of cardiac troponin-I (cTNI) measured in serum of WT and cMKO mice at 18 [WT (n=6) vs cMKO (n=6)] and 34 weeks [WT (n=11) vs cMKO (n=20)]. Data represent mean ± SD; 2-tailed unpaired Student’s t-test at 18 weeks and 34 weeks (w); ***p<0.001. **O) M**yosin light chain 1 (MLC-1) levels measured in serum of WT and cMKO mice at 18 [WT (n=5) vs cMKO (n=6)] and 34 weeks (w) [WT (n=11) vs cMKO (n=17)]. Data represent mean ± SD; 2-tailed unpaired Student’s t-test at 18w and 34w; **p<0.01 and ***p<0.001. **P)** Volcano plots generated from the RNAseq analysis (Dataset EV2) of the differentially expressed genes in cardiac tissue of WT and cMKO mice at 8 (left) and 18 weeks (right). **Q)** Number of genes up-regulated and down-regulated in cMKO mice at 8 (blue) and 18 (pink) weeks (w) within the gene ontology (GO) term: metabolic process (left), mitochondrial genes (MitoCarta, middle) and inflammation (right) obtained from RNAseq analysis (Dataset EV2). **R)** mRNA expression measured by bulk RNAseq (arbitrary units; AU) of the indicated profibrotic genes in heart tissue of WT (n=6) and cMKO (n=6) mice at 18 weeks. Data represent mean ± SD; 2-tailed unpaired Student’s t-test.

To directly assess the importance of MTFP1 for cardiac structure and function, we performed longitudinal echocardiographic (echo) studies in male and female mice (Figure 1G-L, S1F-K). Echo analyses beginning at 10 weeks of age revealed normal cardiac structure and function in cMKO mice. By 18 weeks, however, despite normal cardiac structure we observed a progressive decrease in systolic function, culminating in dilated cardiomyopathy (DCM) and left ventricle (LV) remodeling. By 34 weeks, cMKO mice exhibited all the hallmarks of DCM and heart failure (HF): significantly reduced LV ejection fraction [% EF, WT 60.31 ± 5.4 % *versus* cMKO 25.87± 11.8 %; Figure 1H], thinning of the interventricular septum in systole [IVS (mm), WT 1.23 ± 0.097 *versus* cMKO 0.9 ± 0.092; Figure 1I), posterior wall during systole [PWs (mm), WT 1.39 ± 0.10 *versus* cMKO 0.952 ± 0.17; Figure 1J), dilated LV chamber during the cardiac cycle of diastole [LVDD (mm), WT 4.00 ±0.15 *versus* cMKO 5.02 ± 0.65, Figure 1K] and systole [LVSD (mm), WT 2.72 ± 0.16 *versus* cMKO 4.43 ± 0.84; Figure 1L) with pulmonary congestion (Figure S1L) and increased heart mass at severe HF (Figure S1M).

To determine whether the progressive cardiac contractile dysfunction observed in cMKO mice was caused by primary defects in cardiomyocyte function, we assessed sarcomere length and shortening (Figure S1N-P) coupled to intracellular Ca^2+^ levels [Ca^2+^]_c_ (Figure S1Q-S) in field-stimulated cMKO cardiomyocytes isolated from pre-symptomatic mice (8-10 weeks of age). Diastolic and systolic sarcomere length as well as [Ca^2+^]_c_ transients were similar between WT and cMKO myocytes both at baseline (0.5 Hz stimulation frequency) and during β-adrenergic stimulation (5 Hz stimulation frequency) despite a modest increase of sarcomere shortening of cMKO myocytes under stress conditions and a normal sarcomere re-lengthening at baseline conditions. Together, these data indicate that *Mtfp1* deletion does not impinge upon cardiomyocyte excitation-contraction coupling before the onset of cardiomyopathy in vivo.

Histological analyses of cMKO hearts of mice at HF (34 weeks) confirmed defects in cardiac structure: hematoxylin-eosin (H&E) staining of cardiac cross sections showed LV chamber expansion and myocardial wall thinning, while Masson’s trichrome and Picrosirius Red staining showed disruption of the myofibril architecture by dramatic fibrosis and collagen deposition at DCM (Figure 1M). We also found increased serum levels of cardiac troponin I (cTNI, Figure 1N) and myosin light chain 1 (MLC-1, Figure 1O) in cMKO mice sampled at 18 and 34 weeks, further substantiating the ongoing cardiomyocyte damage and death. We did not observe gender differences in the development of the cardiac dysfunction (Figure S1F-K), highlighting the essential nature of MTFP1 for cardiac structure and function.

### Metabolic and inflammatory gene expression dysregulation in cMKO mice

To gain further insights into the molecular and cellular mechanisms underlying cardiac pathology in cMKO mice, we performed transcriptomic analyses by bulk RNA sequencing (RNAseq) of LVs isolated from WT and cMKO mice at a pre-symptomatic stage (8 weeks of age) and at the onset of DCM (18 weeks). We observed up-regulation of *Nppa* in pre-symptomatic cMKO mice, which was associated to up-regulation of *Nppb* at the onset of DCM, prototypical cardiomyocyte stress-response genes that are activated in response to hemodynamic load (Hohl et al., 2013) and metabolic or contractile abnormalities (Houweling et al., 2005) (Figure 1P). At the pre-symptomatic stage, we observed limited transcriptional remodeling in hearts from cMKO mice with differential expression of only ∼1% of 25815 genes: 137 genes were downregulated, and 122 genes were upregulated in cMKO mice (Dataset EV2, Figure 1P), whereas at 18 weeks, we observed a much broader transcriptional response with 3642 differentially expressed genes in cMKO mice manifesting early signs of DCM (Dataset EV2, Figure 1P).

Functional enrichment analyses performed with g:Profiler (Reimand et al., 2019) and Enrichr (Chen et al., 2013) revealed a number of dysregulated genes involved in various metabolic processes (Figure 1Q, left). Among the downregulated genes with the gene ontology (GO) term metabolic process (Dataset EV2) were genes required for OXPHOS, TCA cycle, fatty acid oxidation and pyruvate metabolism, suggestive of cardiometabolic changes previously observed in mitochondrial models of HF (Hansson et al., 2004; Liao et al., 2015; Wai et al., 2015; Zhou and Tian, 2018). In fact, further examination revealed that half of all mitochondrial genes referenced on MitoCarta (Rath et al., 2021) were downregulated (Figure 1Q, middle). On the other hand, we observed a strong sterile inflammatory gene expression signature and innate immune engagement (Figure 1Q, right), which together with dysregulated extracellular matrix-remodeling genes [profibrotic cytokines such as TGFb, collagen precursor genes (*Col11a1, Col14a1, Col8a1*) and matrix metalloproteinases (*Mmp-2, Mmp-14, Mmp-3*)] (Figure 1R) corroborates the cardiac fibrosis revealed by histological analysis of cMKO hearts (Figure 1M). Notably, the suppression of mitochondrial gene expression and the activation of sterile inflammation measured in cMKO mice at 18 weeks was absent in pre-symptomatic cMKO mice, implying that these transcriptional responses are downstream consequences *Mtfp1* deletion in adult cardiomyocytes.

### Mtfp1 is required for bioenergetic efficiency in cardiac mitochondria

To directly assess the effects of energy metabolism in cMKO mice, we measured mitochondrial respiration in cardiac mitochondria from WT and cMKO mice (Figure 2A). High resolution respirometry studies showed a general impairment of mitochondrial O_2_ consumption in cMKO hearts at both early and late stage of DCM: complex I-, complex II-and complex IV-driven mitochondrial respiration were all significantly lower at either age (Figure 2B, C). Interestingly, reduced mitochondrial respiration was not accompanied by reduced levels of mitochondrial proteins, including those involved in OXPHOS (Figure S2A-B, Dataset EV1), demonstrating that bioenergetic decline was not the result of increased wholesale mitophagy.

**Figure 2.**
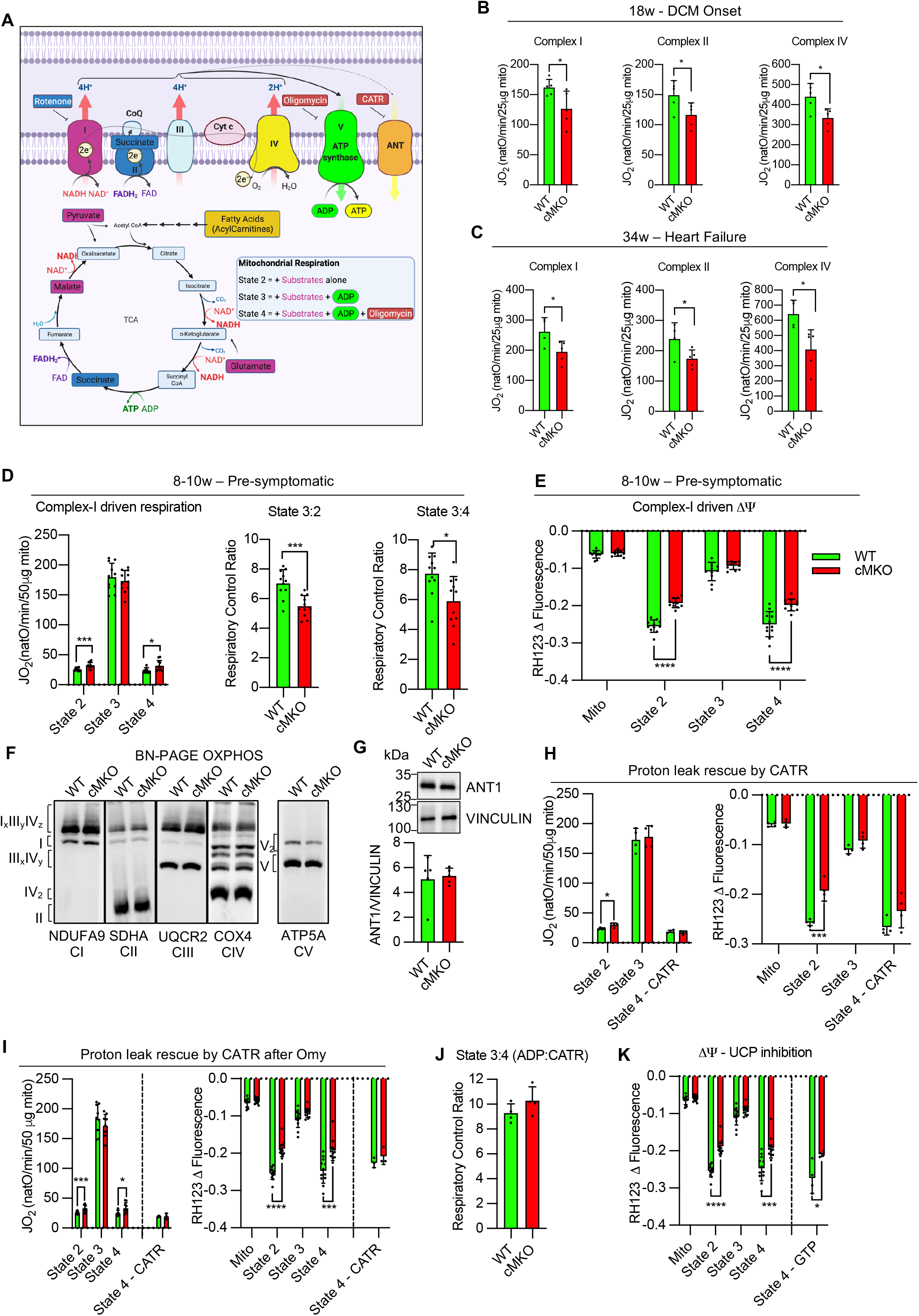
Mtfp1 is required for bioenergetic efficiency in cardiac mitochondria. **A)** Substrates from fatty acid oxidation (mustard) and glycolysis (purple, blue) are metabolized in the TCA cycle which delivers fuels the electron transport chain (ETC) complexes located in the inner mitochondrial membrane by providing NADH and FADH to complexes I (purple) and II (blue), respectively. Complexes I, III and IV extrude protons from matrix into the intermembranous space creating an electrochemical gradient driving the phosphorylation of ADP at the ATP synthase (complex V). The electron flow is limited by the availability of oxygen, a terminal acceptor of the energy depleted electron at the complex IV (cytochrome oxidase). Uncoupling proteins such as ANT promote a proton leak, playing an important role in regulation of membrane potential and oxidative phosphorylation efficiency. Specific inhibitors of complex I (rotenone), complex V (oligomycin), and ANT (carboxyatractyloside, CATR). **B)** Oxygen consumption rates (JO_2_) measured by high-resolution respirometry of cardiac mitochondria from WT (n=5) and cMKO (n=5) male mice at 18 weeks. Complex I driven respiration (left) was measured in presence of pyruvate, malate, glutamate (PGM) and ADP followed by the addition of rotenone and succinate to assess complex II driven respiration (middle) and antimycin A, carbonyl cyanide m-chlorophenyl hydrazine (CCCP), and N,N,N′,N′-Tetramethyl-p-phenylenediamine (TMPD) and ascorbate to measure complex IV driven respiration (right). Data represent mean ± SD; unpaired Student’s t-test, *p<0.05. **C)** Oxygen consumption rates (JO_2_) measured by high-resolution respirometry of cardiac mitochondria from WT (n=5) and cMKO (n=5) male mice at 34 weeks (w). Complex I driven respiration (left) was measured in presence of pyruvate, malate, glutamate (PGM) and ADP followed by the addition of rotenone and succinate to assess complex II driven respiration (middle) and antimycin A, carbonyl cyanide m-chlorophenyl hydrazine (CCCP), and N,N,N′,N′-Tetramethyl-p-phenylenediamine (TMPD) and ascorbate to measure complex IV driven respiration (right). Data represent mean ± SD; unpaired Student’s t-test, *p<0.05. **D)** Oxygen consumption rates (JO_2_) measured by high-resolution respirometry of cardiac mitochondria isolated from WT (n=11) and cMKO (n=11) mice between 8-10 weeks (left). Respiration was measured in presence of pyruvate, malate, and glutamate (PGM) (state 2) followed by the addition of ADP (state 3) and Oligomycin (Omy-state 4). Data represent mean ± SD. Multiple t-test, state 2 ***p<0.001, state 4 *p<0.05. Respiratory control ratios (RCR) of state 3:2 (middle: JO_2_ ADP/PGM) and respiratory control ratios (RCR) of state 3:4 (right: JO_2_ ADP/Omy). Data represent mean ± SD; 2-tailed unpaired Student’s t-test, ***p<0.001. **E)** Mitochondrial membrane potential (ΔΨ) measured by quenching of Rhodamine 123 (RH123) fluorescence in cardiac mitochondria isolated from WT (n=11) and cMKO (n=11) mice between 8-10 weeks. ΔΨ was measured in presence of pyruvate, malate, and glutamate (PGM) (state 2) followed by the addition of ADP (state 3) and Oligomycin (state 4). Data represent mean ± SD; Multiple t-test, ****p<0.0001. **F)** Representative BN-PAGE immunoblot analysis of cardiac OXPHOS complexes isolated from WT and cMKO mice at 8-10 weeks using the indicated antibodies. **G)** Equal amounts of protein extracted from WT (n=5) and cMKO (n=5) hearts between 8-10 weeks were separated by SDS–PAGE and immunoblotted with the indicated antibodies and quantified by densitometry using VINCULIN as a loading control. Data represent mean ± SD. **H)** Oxygen consumption rates measured by high-resolution respirometry (left; JO_2_) and mitochondrial membrane potential (right; ΔΨ) measured by quenching of Rhodamine 123 (RH123) fluorescence in cardiac mitochondria of WT (n=4) and cMKO (n=4) mice between 8-10 week of age. JO_2_ and ΔΨ were measured in presence of pyruvate, malate, and glutamate (state 2) followed by the addition of ADP (state 3) and carboxyatractyloside (CATR) (state 4). Data represent mean ± SD; Multiple t-test, ***p<0.001. **I)** Oxygen consumption rates measured by high-resolution respirometry (left; JO_2_) and mitochondrial membrane potential (right; ΔΨ) measured by quenching of Rhodamine 123 (RH123) fluorescence in cardiac mitochondria of WT and cMKO mice between 8-10 week of age. Respiration was measured in presence of pyruvate, malate, and glutamate (PGM) (state 2) followed by the addition of ADP (state 3), Oligomycin (state 4) and carboxyatractyloside (state 4-CATR) [WT (n=3) cMKO (n=3)]. Data represent mean ± SD. Multiple t-test, state 2 ***p<0.001, state 4 *p<0.05. **J)** Respiratory control ratio (RCR) of state 3:4 (ADP/CATR) between WT (n=4) and cMKO (n=4) calculated from H). Data represent mean ± SD. **K)** Mitochondrial membrane potential (ΔΨ) measured by quenching of Rhodamine 123 (RH123) fluorescence in cardiac mitochondria of WT and cMKO mice between 8-10 week of age. ΔΨ was measured in presence of pyruvate, malate, and glutamate (state 2) followed by the addition of ADP (state 3), oligomycin (state 4) and GTP (state 4 - GTP) [WT (n=3), cMKO (n=3)]. Data represent mean ± SD; Multiple t-test, *p<0.05, ***p<0.001, ****p<0.0001.

Since reduced mitochondrial content and/or OXPHOS activity has been observed in other genetic models of cardiomyopathy (Burke et al., 2016), we wondered whether impaired mitochondrial respiration observed in failing cMKO hearts reflected the consequences of cardiac dysfunction and cardiac remodeling, rather than the ablation of an essential component or regulator of OXPHOS. We observed no deficits of the mitochondrial Krebs cycle in intact field-stimulated cMKO cardiomyocytes assessed by determining the autofluorescence of NAD(P)H/NAD(P)^+^ and FADH_2_/FAD, intrinsic biomarkers of mitochondrial metabolic activity. NAD(P)H/FAD redox state was monitored during a protocol in which cells were exposed to an increase in stimulation frequency and β-adrenergic stimulation(Bertero et al.), creating a typical ADP-induced oxidation (“undershoot”) followed by Ca^2+^-dependent regeneration (“recovery”) behavior (Cortassa et al., 2003)(Figure S2C). This behavior was similar between cMKO and WT myocytes (Figure S2C), ruling out gross alterations in Krebs cycle activity and Ca^2+^-induced redox adaptation.

To uncover the functional alterations of OXPHOS that potentiate the development of DCM, we performed bioenergetic measurements in cardiac mitochondria isolated from pre-symptomatic mice. Using a dual respirometer-fluorimeter (O2k, Oroboros), simultaneous kinetic measurements of oxygen consumption rates (JO_2_) and mitochondrial membrane potential (ΔΨ) were performed in cardiac mitochondria isolated from pre-symptomatic WT and cMKO mice (8-10 weeks). Active mitochondria were pre-labeled with the potentiometric dye Rhodamine-123 (RH-123) and energized with substrates whose metabolism promotes complex I-[state 2; pyruvate, glutamate, and malate (PGM)] or complex II-linked respiration (state 2; succinate and rotenone), followed by ADP to promote phosphorylating (state 3: ADP) respiration. Finally, the ATP synthase inhibitor oligomycin (Omy) was applied to assess non-phosphorylating (state 4) respiration (Figure 2A).

We observed no differences in state 3 JO_2_ rates between WT and cMKO cardiac mitochondria fueled with either PGM, succinate with rotenone or palmitoyl-carnitine with malate, respectively (Figure 2D (left), Figure S2D (left), F). However, respiration was ∼30% higher in cMKO mitochondria in state 2 (+27.4 %) and state 4 (+30.3 %) when complex I was fueled with PGM (Figure 2D), significantly decreasing respiratory control ratios [State 3:2, State 3:4, Figure 2D (middle, right)]. In addition, altered JO_2_ rates in cMKO mitochondria were accompanied by impaired RH-123 quenching in state 2 and state 4 (Figure 2E, S2E), indicating defective IMM substrate-dependent hyperpolarization. Supporting the notion that MTFP1 ablation increases proton leak, we consistently observed reduced respiratory control ratios when complex II was energized (Figure S2D, middle-right) and lower mitochondrial membrane potential under both state 2 and state 4 conditions regardless of the respiratory substrates that were provided (Figure S2D-E).

The elevated JO_2_ rates and reduced RH-123 quenching in state 4 could be explained either by a reduced sensitivity of the ATP synthase to Omy treatment or by uncoupling caused by proton leak across the IMM. In the mouse heart, reduced Omy sensitivity can result from defects in the assembly of the ATP synthase that alter the affinity of Omy binding to Complex V via OSCP (Mourier et al., 2014). However, BN-PAGE analyses of cardiac mitochondria isolated from pre-symptomatic cMKO mice revealed no defects in ATP synthase assembly/maintenance (Figure 2F), leaving us with increased proton leak as the most parsimonious explanation for the observed state 4 respiration and membrane potential differences (Figure 2D, E; Figure S2D-E). Next, we sought to corroborate our findings in cultured cells. To this end, we generated MTFP1-deficient mouse embryonic fibroblasts (MEFs) (*Mtfp1*^-/-^) and MTFP1-deficient human U2OS cells by Crispr/Cas9 genome editing (*MTFP1^Crispr^*) and corresponding WT (*MTFP1^+/+^*) controls (Figure S2H, I) and then assessed oxygen consumption by Seahorse FluxAnalyzer. Intriguingly, we observed no changes in basal or maximal respiration rates nor any evidence of mitochondrial uncoupling (Figure S2J-Q) suggesting that MTFP1-dependent proton leak may be cell type specific.

Taken together, our data reveal an unappreciated and critical role of MTFP1 in bioenergetic efficiency and mitochondrial uncoupling, particularly evident in metabolically active cardiomyocytes, which precedes the manifestation of cardiac dysfunction and heart failure in cMKO mice.

### Mitochondrial uncoupling is mediated by the Adenine Nucleotide Translocase

To uncover the mechanism responsible for the mild mitochondrial uncoupling caused by *Mtfp1* deletion in cardiomyocytes, we turned our attention to known uncoupling proteins. Uncoupling proteins (UCPs, UCP 1/2/3) and adenine nucleotide translocase (ANT) IMM proteins have been reported to be the two main catalysts of futile proton leak in mammalian mitochondria (Brand et al., 2004; Echtay et al., 2002). UCP1 is a bona fide uncoupler that is primarily expressed in brown adipose tissue (Adams, 2000) and shares significant sequence similarity with UCP2 and 3, which are expressed in other tissues (Woyda-Ploszczyca and Jarmuszkiewicz, 2014). ANT is an integral IMM transporter that imports ADP and concomitantly exports ATP between the mitochondrial matrix and IMS (Ruprecht and Kunji, 2021). ANT exists in four different tissue specific isoforms (ANT1, 2, 3, and 4), with ANT1 being the most abundant protein in mitochondria (Lu YW et al, 2017, Brand MD et al, 2005; Karch et al., 2019; JE Kokoszka et al., 2016). ANT1 has long been known for its namesake role as a nucleotide translocator, and recent studies have proven it to be an essential transporter of protons across the IMM in mammals (Bertholet et al., 2019) and a rate-liming factor for proton leak in *Drosophila* (Brand et al., 2005). While the steady state levels of ANT in cardiac tissue of WT and cMKO mice were identical (Figure 2G), we nevertheless sought to functionally assess the contribution of ANT to MTFP1-dependent proton leak in freshly isolated cardiac mitochondria by using the ANT antagonist carboxyatractyloside (CATR), which binds irreversibly to ANT on the IMS side of the IMM, blocking its activity (Todisco et al., 2016). The addition of CATR alone or after Omy treatment rescued the reduced ΔΨ and elevated JO_2_ rates back to WT levels (Figure 2H-I), normalizing the respiratory control ratio for state 3:4 (Figure 2J), suggesting that *Mtfp1* deletion increases ANT-dependent proton leak.

To exclude the unlikely possibility that other uncoupling proteins such as UCPs might contribute to increased proton leak caused by *Mtfp1* ablation, we measured ΔΨ of cardiac mitochondria in the presence of GTP, a pyrimidine nucleotide previously demonstrated to potently inhibit uncoupling in vitro (Macher et al., 2018; Woyda-Ploszczyca and Jarmuszkiewicz, 2014). UCP inhibition with GTP was not able to rescue the defective membrane potential observed in Omy-treated cMKO mitochondria (Figure 2K) suggesting that UCPs do not contribute to the futile proton leak in cMKO cardiac mitochondria. Taken together, these data demonstrate that depletion of MTFP1 in the IMM leads to an increased uncoupling activity of ANT, resulting in proton leak and bioenergetic inefficiency preceding the development of cardiac dysfunction (Figure S2R).

### MTFP1 is dispensable for mitochondrial fission

Transient MTFP1 knock-down has previously been reported to promote mitochondrial elongation in a variety of cultured cell types including neonatal cardiomyocytes (Duroux-Richard et al., 2016; Morita and Terada, 2015; Tondera et al., 2009; Wang et al., 2017), yet the consequence in the adult heart has never been explored. Therefore, to determine the impact of *Mtfp1* deletion on mitochondrial morphology, we co-labeled primary WT and cMKO primary adult cardiomyocytes (CMs) with tetramethylrhodamine ethyl ester (TMRE) and Mitotracker Deep Red (MTDR) to visualize mitochondria. Contrary to previous reports purporting that *Mtfp1* depletion causes mitochondrial elongation (Aung et al., 2017a, 2019), we failed to observe any obvious effects on the morphology, distribution, or content of mitochondria in primary adult CMs deleted of *Mtfp1* (Figure 3A-B). Similarly, transmission electron microscopy (TEM) analyses of pre-symptomatic cMKO and WT hearts (LV) showed no indications of mitochondrial elongation (median mitochondrial surface area: WT 3198 μm^2^ versus cMKO 2954 μm^2^) nor altered cristae organization (Figure 3C-D). We also did not observe changes in mtDNA content (Figure 3E) or mitochondrial shaping proteins in cardiac biopsies: the steady-state levels of fusion (MFN1, MFN2, OPA1) and fission (DRP1, MID51, FIS1) proteins were no different between WT and cMKO mice (Figure 3F). Thus, MTFP1 is dispensable for mitochondrial dynamics in the heart.

**Figure 3.**
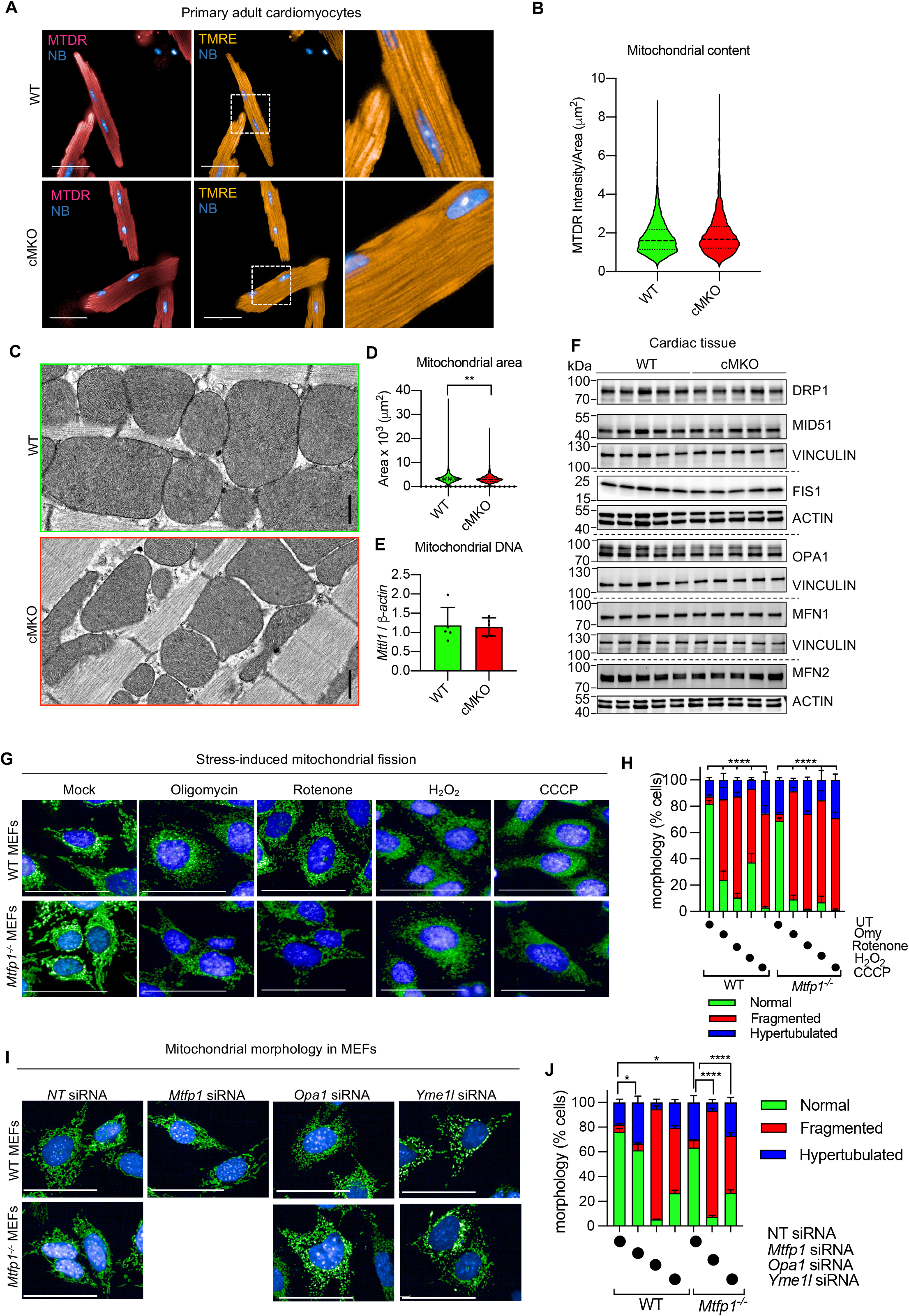
MTFP1 is dispensable for mitochondrial fission. **A)** Representative confocal images of primary adult cardiomyocytes isolated from WT and cMKO mice at 8 weeks. Mitochondria were stained with MitoTracker Deep Red (MTDR) and tetramethylrhodamine, ethyl ester (TMRE), nuclei were stained with NucBlue (NB). Scale bar=50 μm. **B)** Violin plot of mitochondrial content (MTDR Intensity/Area) of WT (n=6085) and cMKO CMs (n=3647) measured in A). **C)** Representative transmission electron micrographs of cardiac posterior walls of WT (top, n=3) and cMKO (bottom, n=3) mice at 8-10 weeks. Scale bar: 500 nm. **D)** Violin plot of mitochondrial surface area (μm^2^) within cardiac posterior wall measured in C (WT mitochondria n=659; cMKO mitochondria n=966). Dotted line represents quartiles and dashed line represents median; **p<0.01 Mann-Whitney test. **E)** Mitochondrial DNA (mtDNA) content in WT (n=5) and cMKO (n=5) heart tissue quantified by amplification of the mitochondrial *Mttl1* gene relative to nuclear gene *b-Actin*. Data represent mean ± SD. **F)** Immunoblot of mitochondrial fission and fusion proteins. Equal amounts of protein from cardiac WT and cMKO (8-10 week) extracts were separated by SDS–PAGE and immunoblotted with the indicated antibodies (horizontal line denotes different membranes). VINCULIN or ACTIN are used as loading controls. **G)** Representative confocal images of WT and *Mtfp1^-/-^* MEFs treated with the fission inducing drugs: oligomycin (Omy), Rotenone, H_2_O_2_ and carbonyl cyanide m-chlorophenyl hydrazine (CCCP). Mitochondria were stained with MitoTracker Deep Red (MTDR, green) and nuclei with NucBlue (NB, blue). Scale bar=100 μm. **H)** Quantification of mitochondrial morphology in G) by supervised ML using WT cells with normal (UT), fragmented (CCCP-treated) or hypertubular (cycloheximide-treated) mitochondria as ground truths. Data are means ± SEM of 4 independent replicates. 2way-ANOVA, Dunnet’s multiple comparison test: % fragmentation ****p<0.0001 treatment vs WT UT or *Mtfp1^-/-^* UT. **I)** Representative confocal images of WT and *Mtfp1^-/-^* MEFs treated with indicated siRNAs (20 nM) for 72 h. Mitochondria were stained with MitoTracker Deep Red (MTDR, green) and nuclei with NucBlue (NB, blue). Scale bar=100 μm. **J)** Quantification of mitochondrial morphology in I) by supervised machine learning (ML) using WT cells with normal (non-targeting NT siRNA), fragmented (*Opa1* siRNA) or hypertubular (*Dnm1l* siRNA) mitochondria as ground truths. Data are means ± SEM of 4-8 technical replicates. 2way-ANOVA, Dunnet’s multiple comparison test: % hypertubular * p<0.05; ***p<0.001 versus WT NT siRNA; % fragmented ****p<0.0001 versus *Mtfp1^-/-^* NT siRNA.

In light of these surprising findings, we decided to measure mitochondrial morphology in MEFs depleted of MTFP1 under basal and stress conditions using a recent supervised machine learning (ML) approach we developed for high-throughput image acquisition and analyses (Cretin et al., 2021). In contrast to previous reports, MEFs depleted (siRNA *Mtfp1*) or deleted (*Mtfp1^-/-^)* of *Mtfp1* showed only modest elongation of the mitochondrial network: both transient (siRNA) or chronic (knockout) ablation of MTFP1 resulted in ∼15% increase in hypertubular mitochondria (Figure 3I-J), and *Mtfp1^-/-^* MEFs showed unaltered steady-state levels of mitochondrial fusion and fission proteins (Figure S3A). Contrary to DRP1-deficient cells (Cretin et al., 2021), *Mtfp1^-/-^* MEFs were not protected from mitochondrial fragmentation induced by established pharmacological triggers of mitochondrial fragmentation, such as oligomycin (Omy), Rotenone, hydrogen peroxide (H_2_O_2_), or carbonyl cyanide m-chlorophenylhydrazone (CCCP) (Figure 3G-H). Similarly, *Mtfp1^-/-^* MEFs were not protected from genetic induction of mitochondrial fragmentation by depletion of *Yme1l* or *Opa1* (Figure 3I-J). While we could confirm that MTFP1 overexpression is able to promote mitochondrial fragmentation in MEFs, without affecting steady state level of fusion and fission proteins (Figure S3B-C), our data collectively indicate that MTFP1, unlike DRP1, is not an essential fission protein. Taken together, these results strongly argue that contrary to its namesake, MTFP1 is not an essential fission factor either in vitro or in vivo.

### MTFP1 deletion promotes mitochondrial permeability transition pore opening and programmed cell death

Cardiomyocyte death is catastrophic for adult cardiac function because of the limited regenerative capacity of these post-mitotic cells. Given the appearance of cardiac cell damage and death in cMKO mice during DCM (Figure 1N-O), we sought investigate whether MTFP1 loss specifically increased cell death sensitivity. To this end, we isolated adult primary cardiac myocytes (CMs) from WT and pre-symptomatic cMKO mice between 8-10 weeks of age and kinetically measured cell survival in response to a variety of cell death triggers using supervised ML-assisted high-throughput live-cell imaging (Cretin et al., 2021). We were able to isolate equally viable CMs from both WT and cMKO mice, yet upon dissipation of the membrane potential with CCCP (Figure 4A-B) or treatment with H_2_O_2_ (Figure 4C-D), *Mtfp1^-/-^* CMs succumbed to cell death more rapidly than WT CMs. Moreover, the induction of cell death with doxorubicin (DOXO, Figure 4E-F), a cardiotoxic chemotherapeutic agent that triggers programmed cell death (PCD) (Christidi and Brunham, 2021), induced a significant increase of death in *Mtfp1^-/-^* compared to WT CMs, indicating that MTFP1 is essential for cell survival.

**Figure 4.**
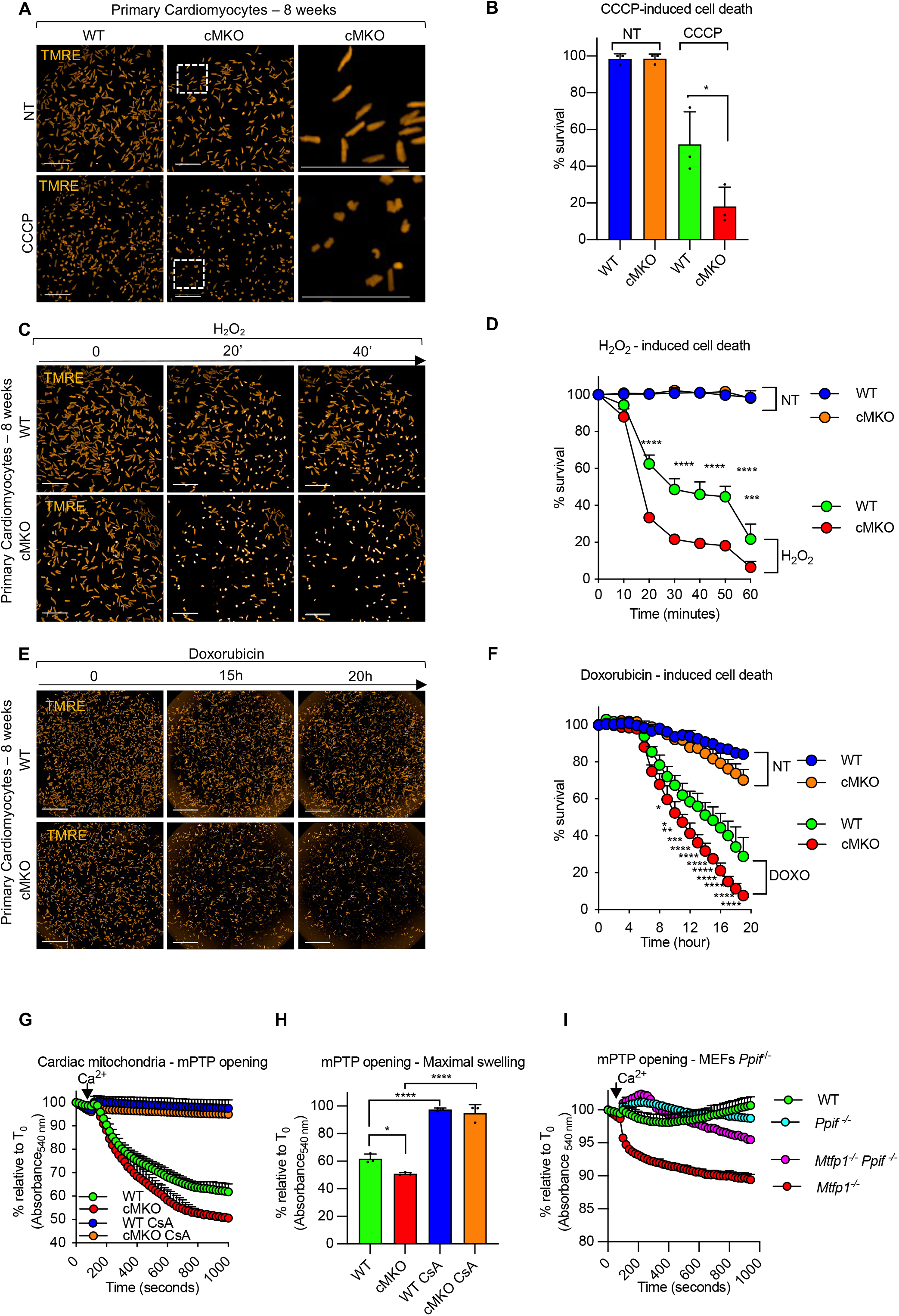
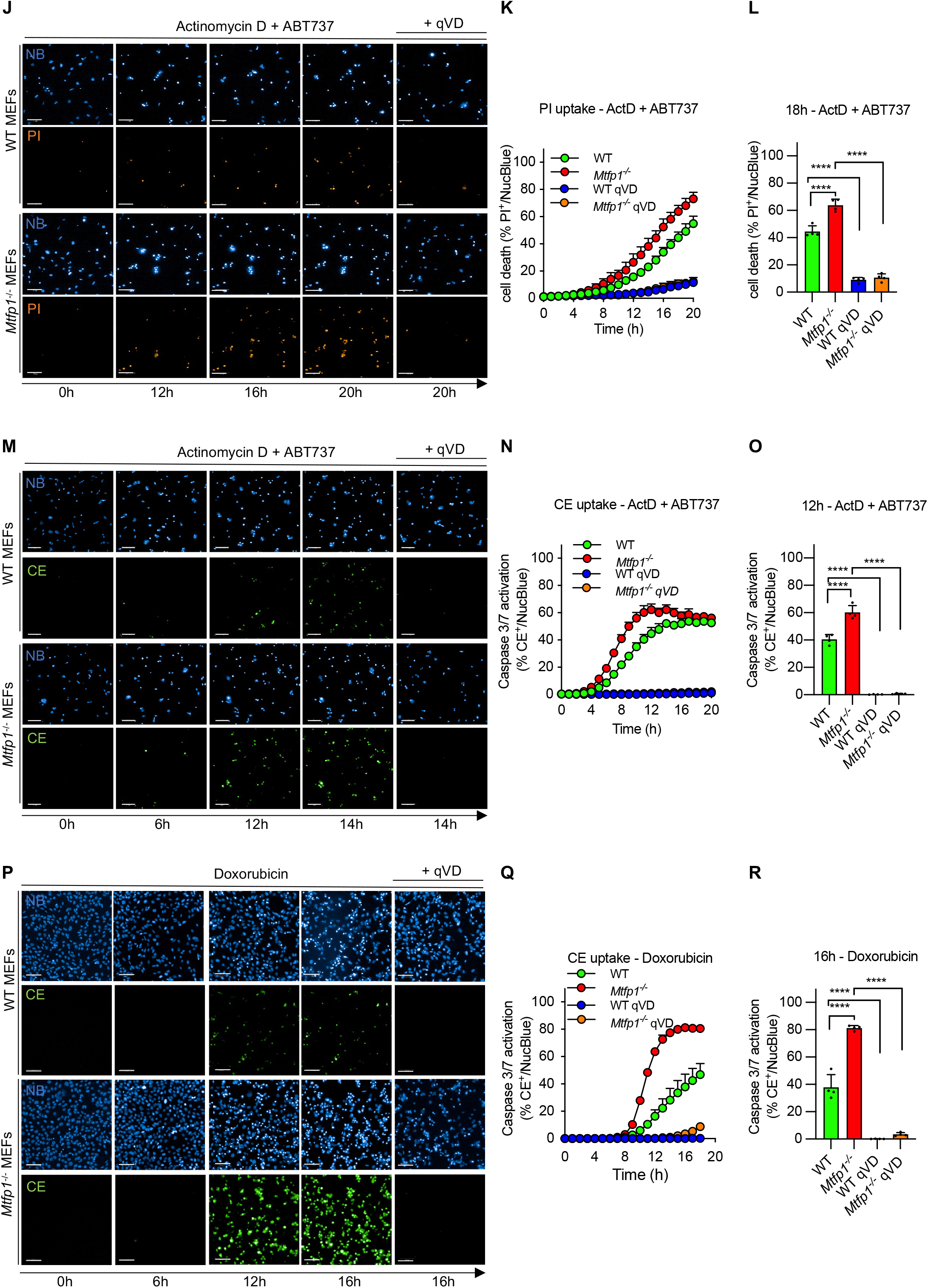
MTFP1 protects against mitochondrial PTP opening and cell death. **A)** Representative confocal images of adult cardiomyocytes (CMs) isolated from WT and cMKO at 8 weeks stained with tetramethylrhodamine, ethyl ester (TMRE) treated with or without cyanide m-chlorophenyl hydrazine treatment (CCCP) for 15 minutes. Rod-shaped CMs: live cells, round-shaped CMs: dead cells. Scale bar: 500 μm. **B)** Quantification of number of live cells (% survival) in A) by supervised machine learning. Data are means ± SD of n=3 independent experiments. Unpaired Student’s t-test; *p<0.05. **C)** Representative confocal images of adult cardiomyocytes (CMs) isolated from WT and cMKO at 8 weeks stained with tetramethylrhodamine, ethyl ester (TMRE) and subjected to H_2_O_2_ treatment for 1 hour. Rod-shaped CMs: live cells, round-shaped CMs: dead cells. Scale bar: 500 μm. **D)** Quantification of number of live cells (% survival) over time measured in C) by supervised machine learning. Data are means ± SD of 2-3 culture replicates and representative of n=3 experiments. 2wayANOVA, Tukey’s multiple comparison test, ***p<0.001, ****p<0.0001 vs WT H_2_O_2_. **E)** Representative confocal images of adult cardiomyocytes (CMs) isolated from WT and cMKO at 8 weeks stained with tetramethylrhodamine, ethyl ester (TMRE) and subjected to Doxorubicin (DOXO). Scale bar: 500 μm. **F)** Quantification of number of live cells (% survival) in E) by supervised machine learning. Data are means ± SD of 2-3 culture replicates and representative of n=3 experiments. 2wayANOVA, Tukey’s multiple comparison test, *p<0.05, **p<0.01, ***p<0.001, ****p<0.0001 vs WT DOXO. Scale bar: 1 mm. **G)** Mitochondrial permeability transition pore (mPTP) opening. Swelling assay performed on cardiac mitochondria extracted from hearts of WT (n=3) and cMKO (n=3) mice at 8-10 weeks. Light scattering (absorbance 540 nm, % relative to T_0_] was measured every 20 s before and after addition of a single pulse of CaCl_2_ (arrowhead). Cyclosporin A (CsA) was added prior CaCl_2_ pulse to inhibit mPTP-induced mitochondrial swelling. Data are means ± SD. **H)** one-way ANOVA of maximal absorbance_540 nm_ (% relative to T_0_) change in G) *p<0.05, ****p<0.0001. **I)** Mitochondrial permeability transition pore (mPTP) opening. Swelling assay performed on mitochondria purified from WT and *Mtfp1^-/-^, Ppif^-/-^* and *Mtfp1^-/-^Ppif^-/-^* MEFs. Mitochondrial absorbance changes (absorbance 540 nm, % relative to T_0_) are measured every 20 s prior and after addition of a single pulse of CaCl_2_ (arrowhead). Cyclosporin A (CsA) was added prior Ca^2+^ to inhibit mPTP-dependent swelling. Data are means ± SD. **J)** Representative confocal images of WT and *Mtfp1^-/-^* MEFs subjected to actinomycin D (ActD) *plus* ABT-737 treatment in the presence or absence of the pan-caspase inhibitor q-VD-OPh hydrate (qVD). Cell death was monitored by using nuclear Propidium Iodide uptake (PI, orange) and imaging cells every hour (h) for 20h. Scale bar = 100 μm. **K)** Kinetics of PI uptake was determined by counting the number of PI^+^ positive cells (orange) over total number cells nuclear stained with NucBlue (NB, blue) and expressed as % PI^+^/NucBlue. Data are means ± SD of n=4 independent experiments. **L)** one-way ANOVA of K) at 18h, ****p<0.0001. **M)** Representative confocal images of WT and *Mtfp1^-/-^* MEFs subjected to actinomycin D (ActD) plus ABT-737 treatment in the presence or absence of the pan-caspase inhibitor q-VD-OPh hydrate (qVD). Live induction of the caspase 3/7 activation was monitored by using the CellEvent (CE, green) reagent and imaging cells every hour (h) for 20h. Scale bar = 100 μm. **N)** Kinetics of caspase 3/7 activation was determined by counting the number of CE^+^ positive cells (green) over total number cells nuclear stained with NucBlue (NB, blue) and expressed as % CE^+^/NucBlue. Data are means ± SD of n=4 independent experiments. **O)** one-way ANOVA of N) at 12h, ****p<0.0001. **P)** Representative confocal images of WT and *Mtfp1^-/-^* MEFs subjected to doxorubicin treatment in the presence or absence of the pan-caspase inhibitor q-VD-OPh hydrate (qVD). Live induction of the caspase 3/7 activation was monitored by using the CellEvent (CE, green) reagent and imaging cells every hour (h) for 18h. Scale bar = 100 μm. **Q)** Kinetics of caspase 3/7 activation was determined by counting the number of CE^+^ positive cells (green) over total number cells nuclear stained with NucBlue (NB, blue) and expressed as % CE^+^/NucBlue. Data are means ± SD and representative of at least n=3 independent experiments. **R)** one-way ANOVA of Q) at 12h, p<0.0001.

Prolonged opening of mitochondrial permeability transition pore (mPTP) causes mitochondria swelling, membrane potential dissipation and bioenergetic collapse, becoming a determinant of cell death (Carraro et al., 2020). To test whether MTFP1 loss causes increased susceptibility to mPTP opening, we assessed mitochondria swelling by exposing cardiac mitochondria of pre-symptomatic WT and cMKO mice to a high concentration of Ca^2+^ and kinetically measured the light scattering (Karch et al., 2013). Ca^2+^ overload induced an increased mPTP dependent swelling of MTFP1-deficient cardiac mitochondria, which could be inhibited by the mPTP inhibitor cyclosporin A (CsA), indicating that *Mtfp1* deletion sensitizes cardiac mitochondria to mPTP opening (Figure 4G-H). These findings were corroborated in *Mtfp1^-/-^* MEFs, in which we observed increased mPTP sensitivity that could be suppressed by CsA treatment (Figure S4A) or by knocking out Cyclophilin D (Cyp D, encoded by *Ppif*), the pharmacological target of CsA in mitochondria (Figure 4I) (Halestrap and Davidson, 1990). Thus, these data clearly indicate that loss of MTFP1 at the level of the IMM sensitizes mitochondria to mPTP opening. To define the molecular mechanisms underlying the increased sensitivity PCD and mPTP opening caused by MTFP1 ablation, we used WT and *Mtfp1^-/-^* MEFs. We began by confirming that like *Mtfp1^-/-^* CMs, MEFs deleted of *Mtfp1* were more sensitive to PCD. As expected, *Mtfp1^-/-^* MEFs showed increased sensitivity to multiple cell death stimuli, as evidenced by more rapid kinetics of caspase 3/7 activation and cell death monitored by ML-assisted live-cell imaging of CellEvent (CE) and Propodium Iodide (PI) uptake, respectively (Cretin et al., 2021). Treatment with cell death triggers actinomycin D (ActD) and ABT-737 (Figure 4J-O), staurosporine (STS, figure S4B-D) or DOXO (Figure 4P-R) all promoted a more rapid and robust cell death response in *Mtfp1^-/-^* MEFs relative to WT MEFs, which could be blocked with the pan-caspase inhibitor Q-VD-OPh (qVD). These effects were independent of cell proliferation or mitochondrial respiration, neither of which were altered in *Mtfp1^/-^* MEFs (Figure S4E, S2J-M). Taken together, our data clearly demonstrate a protective role of MTFP1 in maintaining cell integrity and survival.

### Doxorubicin induced-cardiotoxicity accelerates the onset of cardiomyopathy in cMKO mice

To test whether MTFP1 protected against PCD induction *in vivo*, we injected pre-symptomatic (aged 8 weeks) cMKO and WT mice with DOXO and assessed cardiac function at 14 days post treatment by echo (Figure S4F-J). Consistent with the data obtained *in vitro*, we observed that DOXO accelerated the onset of cardiac dysfunction in cMKO mice by lowering LVEF (WT 60.85 ± 6.2% vs cMKO 47.02 ± 9.1%), PW thickness during systole (WT 1.082 ± 0.099 mm vs cMKO 0.887 ± 0.062 mm), while increasing systolic LV diameter (WT 2.393 ± 0.27 mm vs cMKO 3.009 ± 0.36 mm) and diastolic LV diameter (WT 3.505 ± 0.20 mm vs cMKO 3.918 ± 0.25 mm). These results clearly indicate that cMKO mice are more susceptible to DOXO induced cardiotoxicity, accelerating the onset of cardiomyopathy.

### Inhibition of mPTP rescues cell death sensitivity of MTFP1 deficient cells

It has been previously reported that DOXO mediates mPTP opening and cell death in lung cancer cells (Lu et al., 2014) and cardiac myocytes, and that H_2_O_2_ activates necrosis through the induction of mPTP opening (Vaseva et al., 2012). We observed that H_2_O_2_-induced cell death was accelerated in *Mtfp1^-/-^* cells and was reduced, yet not totally abolished by caspase inhibition with qVD, indicating that MTFP1 loss also renders cells more susceptible to caspase-independent cell death (Figure 5A-C). Since MTFP1-deficient cardiac and MEF mitochondria were more susceptible to mPTP opening (Figure 4G-I, S4A), we investigated whether prolonged mPTP opening contributes to increased cell death sensitivity of MTFP1-deficient cells. To test the dependence of PCD on Cyp D, we disrupted Cyp D in WT and *Mtfp1^-/-^* MEFs by introducing a truncating, homozygous frame shift mutation (p.Val65*) by Crispr/Cas9 genome editing (Figure 5D) and subjected cells to H_2_O_2_ (Figure 5E-G) or DOXO treatment (Figure 5H-J). By tracking the kinetics of PI or CE uptake respectively, we observed that Cyp D ablation in *Mtfp1^-/-^* cells (*Mtfp1^-/-^Ppif1^-/-^* MEFs) rescued the cell death sensitivity back to WT levels (Figure 5E-J). Consistent with a cytoprotective effect of Cyp D ablation, we observed that the association of CsA to qVD treatment had a synergic effect in suppressing cell death sensitivity of *Mtfp1^-/-^* MEFs to WT levels (Figure 5A-C). Taken together, these results clearly indicate that loss of MTFP1 promotes mPTP opening via Cyp D to lower the resistance to programmed cell death. To gain insights into the molecular regulation of the mPTP by MTFP1 we sought to define the cardiac interactome of MTFP1. We expressed FLAG-MTFP1 at the *Rosa26* locus in C57Bl6/N mouse hearts via targeted transgenesis (Figure S5A) and confirmed that the modest level of over-expression did negatively impact cardiac function in vivo (Figure S5B-C). Next, we performed a co-immunoprecipitation study coupled to mass spectrometry (MS) for the analysis of cardiac MTFP1 interactome. We identified 60 mitochondrial proteins besides the bait protein (MTFP1) that were exclusively present in FLAG-MTFP1 eluates or significantly enriched greater than two-fold (Dataset EV3, Figure S5D). Among these interactors we found factors involved in OXPHOS function (Figure S5E), notably proteins required for the assembly and functions of Complex I (Ndufa10, NDUFA7, NDUFS6, NDUFB4, NDUFS6, Mtnd5), Complex IV (CMC2, COX4I1, COX6B1, COX7A1, SCO2), and Complex V (ATP5L, USMG5). In addition, we identified a number of proteins that have previously been implicated in mPTP regulation including the ADP/ATP translocase SLC25A4 (Karch et al., 2019), the inorganic phosphate carrier SLC25A3 (Kwong et al., 2014), and the heat shock protein TRAP1 (Kang et al., 2007) (Figure S5D-E). Altogether, our data have uncovered a functional and physical link between MTFP1 and the mPTP complex in the inner mitochondrial membrane.

**Figure 5.**
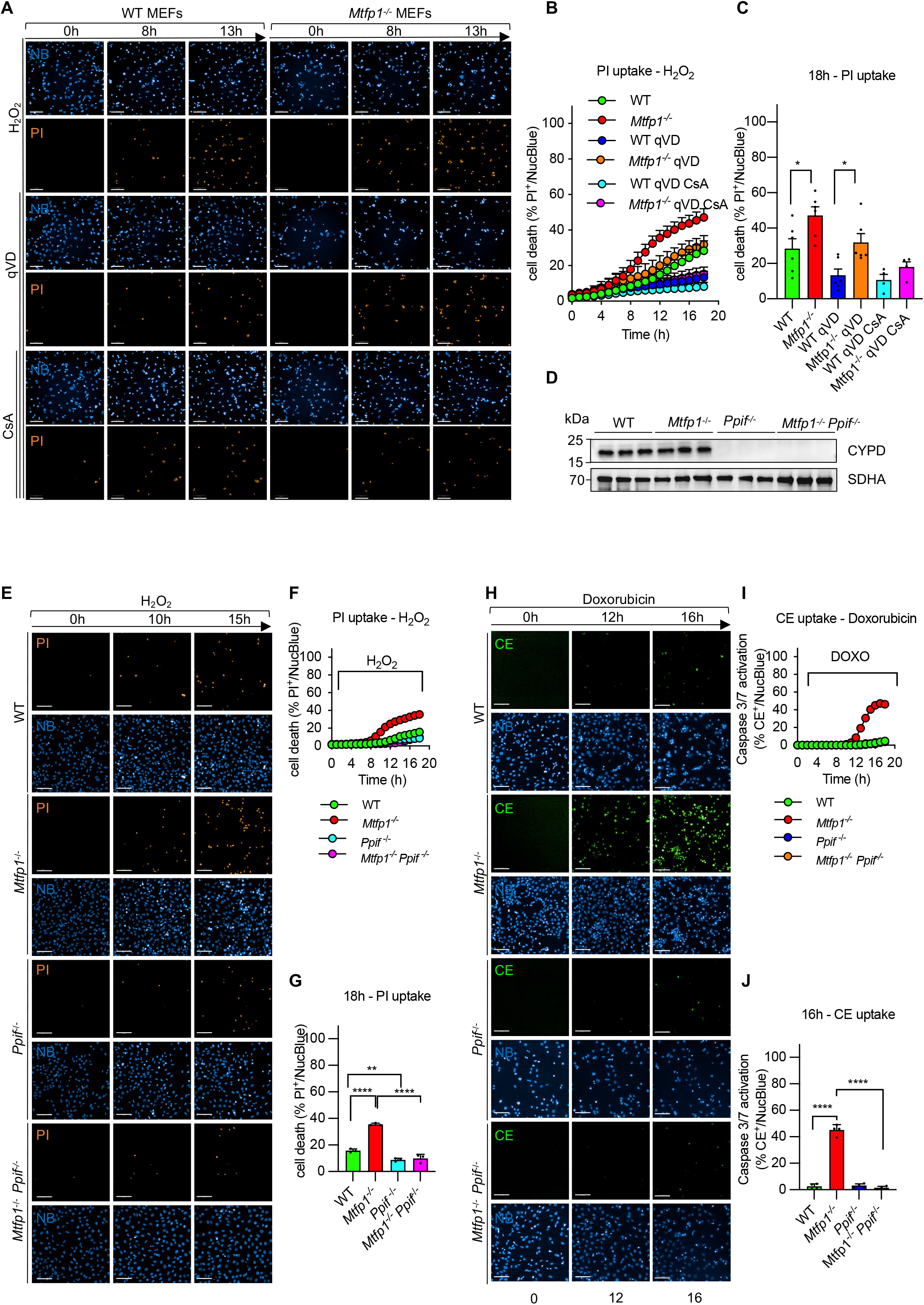
mPTP accounts for cell death sensitivity in MTFP1 deficient cells. **A)** Representative confocal images of WT and *Mtfp1^-/-^* MEFs subjected to H_2_O_2_ treatment. The pan-caspase and cyclophilin D inhibitors, q-VD-OPh hydrate (qVD) and cyclosporin A (CsA) respectively, were used to block both caspase and mPTP dependent cell death. Cell death was monitored by Propidium Iodide uptake (PI, orange) and imaging cells every hour (h) for 18h. Scale bar = 100 μm. **B)** Kinetics of PI uptake was determined by counting the number of PI^+^ positive cells (orange) over total number cells nuclear stained with NucBlue (NB, blue) and expressed as % PI^+^/NucBlue. Data are means ± SD of n=6 independent experiments. **C)** one-way ANOVA of B) at 18h, *p<0.05 **D)** Validation of Cyclophilin D (CYP D) deletion by Crispr/Cas9 genome editing in WT and *Mtfp1^-/-^* MEFs. Equal amounts of protein extracted from WT, *Mtfp1*^-/-^, *Ppif^-/-^*, *Mtfp1*^-/-^ *Ppif^-/-^* MEFs were separated by SDS–PAGE and immunoblotted with the indicated antibodies. SDHA was used as mitochondrial marker and loading control. **E)** Representative confocal images of WT, *Mtfp1*^-/-^, *Ppif^-/-^*, *Mtfp1*^-/-^ *Ppif^-/-^* MEFs subjected to H_2_O_2_ treatment. Cell death was monitored by Propidium Iodide uptake (PI, orange) and imaging cells every hour (h) for 18h. Scale bar = 100 μm. **F)** Kinetics of PI uptake was determined by counting the number of PI^+^ positive cells (orange) over total number cells nuclear stained with NucBlue (NB, blue) and expressed as % PI^+^/NucBlue. Data are means ± SD of n=3 independent experiments. **G)** one-way ANOVA of F) at 18h; **p<0.01, ****p<0.0001. **H)** Representative confocal images of WT, *Mtfp1*^-/-^, *Ppif^-/-^*, *Mtfp1*^-/-^ *Ppif^-/-^* MEFs subjected to doxorubicin (DOXO) treatment. Live induction of the caspase 3/7 activation was monitored by using the CellEvent (CE, green). CellEvent positive cells (CE^+^, green) over total number over cells NucBlue labeled (blue) were imaged every hour (h) for 18h. Scale bar = 100 μm. **I)** Kinetics of caspase 3/7 activation was determined by counting the number of CE^+^ positive cells (green) over total number cells nuclear stained with NucBlue (NB, blue) and expressed as % CE^+^/NucBlue. Data are means ± SD and representative of n=3 independent experiments. **J)** one-way ANOVA of I) at 16h, ****p<0.0001.

**Figure 6.**
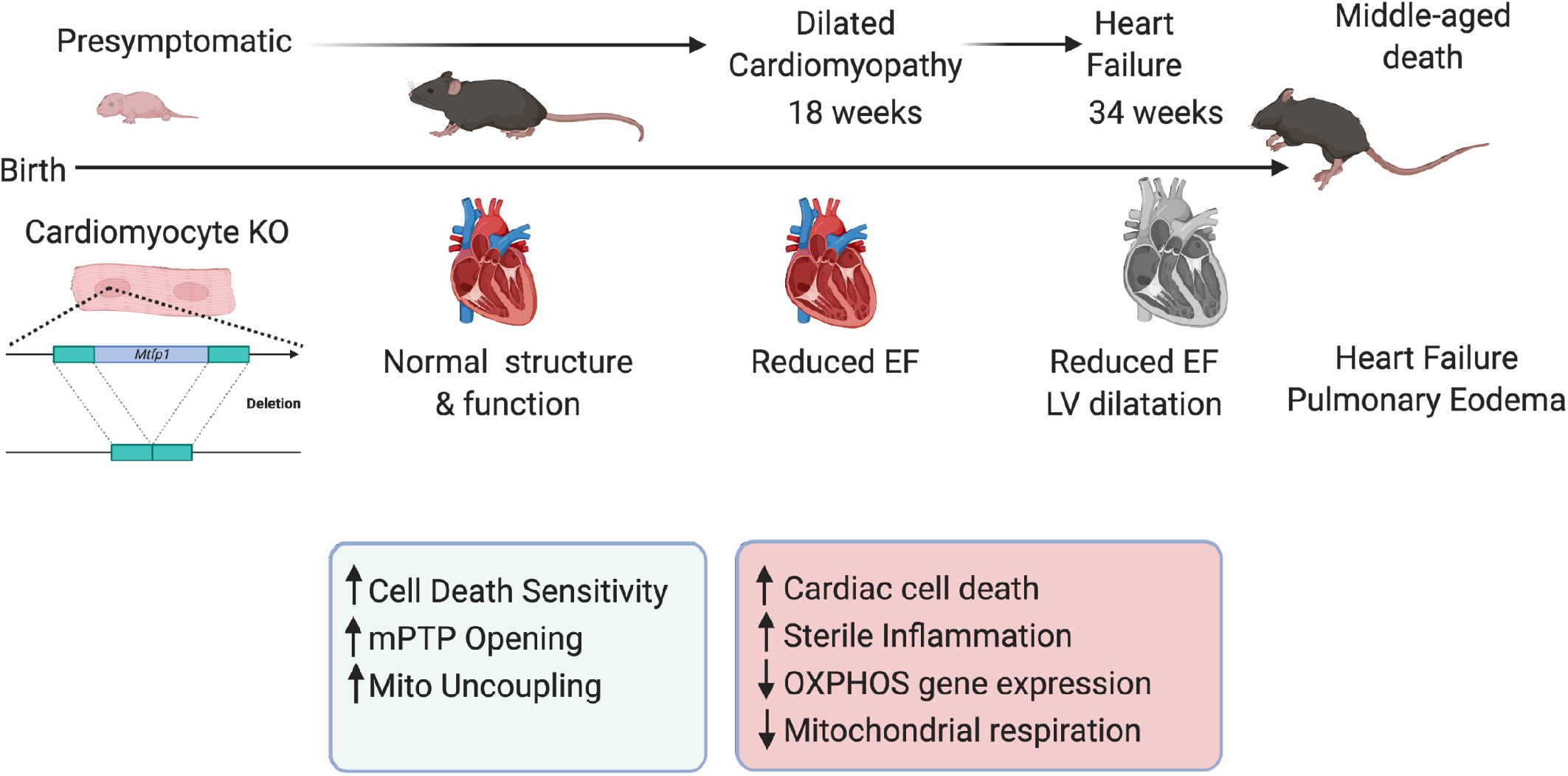
Model for the regulation of mitochondrial and cardiac function by MTFP1. *Mtfp1* deletion in cardiomyocytes occurs at birth (cMKO) and sensitizes cardiac myocytes to mitochondrial permeability transition pore (mPTP) opening, cell death and increases mitochondria uncoupling of the inner membrane. At the adult age of 8-10 weeks heart of cMKO mice have normal structure and function but undergoes to the development of a progressive dilated cardiomyopathy (DCM) at 18 weeks which progresses to severe heart failure and middle-aged death by 34 weeks. At onset of DCM, cMKO mice exhibit increased cardiac cell death, reduced mitochondrial respiration, and induction of a sterile inflammatory response.

## Discussion

We initially chose to focus studies on Mitochondrial Fission Process 1 (MTFP1) because we viewed this protein as a promising entry point to study the hitherto molecularly undefined process of inner membrane division (Wai and Langer, 2016), based largely on previous studies that had purported a pro-fission role upon over-expression and an anti-fission role upon depletion (Aung et al., 2017a, 2017b; Duroux-Richard et al., 2016; Morita et al., 2017; Wang et al., 2017). While we demonstrate MTFP1 to be a bona fide inner membrane protein in vivo (Figure S1B-C), confirming previous in vitro studies (Tondera, 2005; Tondera et al., 2004), careful, unbiased mitochondrial morphology analyses unequivocally excluded this protein as an essential fission factor (Figure 3G-J) for the following reasons: 1) acute or chronic depletion of MTFP1 in vitro (Figure 3G-J) or deletion in vivo (Figure 3C-D) had little impact on the elongation of the mitochondrial network and 2) MTFP1-deficient cells were not protected against mitochondrial fragmentation caused either by accelerated fission or impaired fusion induced either by genetic or pharmacological triggers (Figure 3G-J). Consistent with these findings, we did not identify MTFP1 as a regulator of mitochondrial morphology in a recent, comprehensive siRNA-based phenotypic imaging screens of all mitochondrial genes performed in human fibroblasts (Cretin et al., 2021), which did identify essential fission proteins like DRP1 and receptors (Fonseca et al., 2019; Losón et al., 2013; Osellame et al., 2016), forcing us to reconsider the existing models of mitochondrial fission we (Ng et al., 2021; Wai and Langer, 2016) and others had proposed (Giacomello et al., 2020). In contrast, other studies have reported that chemical inhibitors of mTOR (Morita et al., 2017), PI3K (Tondera et al., 2004), or miRNAs (Duroux-Richard et al., 2016; Wang et al., 2017) can deplete MTFP1 protein levels and thus inhibit mitochondrial fragmentation, although the pleiotropic effects of these molecules and the signaling cascades they regulate make it the interpretation of these data very challenging. However, our in vitro studies confirmed that over-expression of MTFP1 does indeed promote mitochondrial fragmentation, which appears to be independent of stress-induced OPA1 processing and alterations in the levels of the fission and fusion executors (Figure S3B-C). In vivo, stable over-expression in cardiomyocytes does not appear to impact basal cardiac function assessed by echocardiography (Figure S5C), although future studies will be required to determine whether MTFP1 overexpression can impact cardiac physiology under stress conditions.

If MTFP1 is dispensable for fission, what role, if any, does this metazoan-specific factor play in mitochondria? Our in vivo studies revealed that cardiomyocyte-specific deletion of MTFP1 (cMKO) drives the progressive development of dilated cardiomyopathy (DCM) beginning at 18 weeks of age culminating in chronic heart failure and middle-aged death in both male and female mice. Cardiac dysfunction was accompanied by a reduction in mitochondrial gene expression and respiratory chain function, fibrotic remodeling (Figure 1P-R) and a general dysregulation of metabolic genes (Figure 1Q), which are features that have been observed in other cardiomyocyte-specific knockout mouse models of mitochondrial genes (Fernandez-Caggiano et al., 2020; Ghazal et al., 2021; Hansson et al., 2004; Liao et al., 2015; Wai et al., 2015; Zhang et al., 2020b). Similarly, accumulation of pathogenic mutations in mitochondrial DNA have recently been shown to drive mitochondrial dysfunction and sterile inflammation in a number of different tissues including the heart and genetic inhibition of the latter phenotype appears to resolve tissue dysfunction, supporting a pathological role of cardiac inflammation triggered by mitochondrial dysfunction(Lei et al., 2021). In cMKO mice, the aforementioned phenotypes were absent before the onset of DCM, indicating that MTFP1 deletion in perinatal cardiomyocytes does not compromise post-natal cardiac development nor function in mice. Thus, we hypothesize that the metabolic and inflammatory remodeling that accompanies DCM manifests as downstream response to cardiac dysfunction, which we are currently testing. Surprisingly, functional characterization of field-stimulated primary adult cardiomyocytes from pre-symptomatic mice did not reveal defects in contractile capacity, excitation contraction (EC) coupling, or sarcomere integrity (Figure S1N-S), prompting us to search for other homeostatic dysfunctions of cardiomyocytes that could account for the contractile defects of the beating heart caused by MTFP1 deletion. Indeed, characterization of primary cardiomyocytes and cardiac mitochondria from cMKO mice revealed an increased sensitivity to programmed cell death and increased sensitivity to opening of the mitochondrial permeability transition pore (mPTP), respectively (Baines et al., 2005; Carraro et al., 2020; Nakagawa et al., 2005). Accelerated opening of the mPTP has been shown to control cardiomyocyte viability and cardiac function in genetic (Song et al., 2015), infectious (Milduberger et al., 2021), and surgically-induced mouse models of cardiomyopathy (Nakagawa et al., 2005). The identity of the mPTP has been hotly debated (Carraro et al., 2020; Kwong and Molkentin, 2015; Morciano et al.) and a consensus has yet to be achieved regarding its structure and molecular constitution, although compelling evidence from knockout mice and in vitro reconstitution experiments have identified a number of mitochondrial proteins associated with or integrated into the IMM subunits that are required for the efficient opening of the mPTP including subunits of the ATP synthase, cyclophilin D (Cyp D, encoded by the nuclear gene *Ppif*), and IMM carrier proteins of the SLC25 family (Baines et al., 2005; Basso et al., 2008; Bernardi et al., 2015; Carraro et al., 2018, 2020; Karch et al., 2013, 2019; Kwong et al., 2014; Nakagawa et al., 2005), several of which we identified by coimmunoprecipitation studies performed on cardiac mitochondria to physically interact with FLAG-MTFP1, including SLC25A3 (Phosphate carrier), SLC25A4 (ANT1), TRAP1, ATP5L and USMG5 (Figure S5D-E). Complexosome profiling of rat cardiac mitochondria revealed co-migration of MTFP1 with several proteins we identified by coimmunoprecipitation (Heide et al., 2012), including SLC25A3, SLC25A4, and Cyp D, implying that MTFP1 may act as a scaffold in the IMM to participate in the maintenance, assembly or regulation of multi-protein complexes such as the mPTP complex. In MTFP1-deficient cells, inhibiting mPTP opening by pharmacological inhibition with Cyclosporin A (CsA) or genetic deletion of Cyp D rescues the sensitivity to mPTP opening and accelerated cell death. Our discovery that the genetic deletion of *Mtfp1* sensitizes both post-mitotic cardiac cells and mitotic epithelial cells to programmed cell death (PCD) without affecting cardiomyocyte differentiation nor cell proliferation argues for a general role for MTFP1 in cell survival, which is supported by studies in other cell lines (Duroux-Richard et al., 2016; Morita et al., 2017). Why other groups have reported that MTFP1 depletion can protect gastric cancer and cardiomyocyte cell lines from PCD induced by various cell death triggers including doxorubicin is unclear (Aung et al., 2017a, 2017b; Wang et al., 2017). Although our in vivo data demonstrate that MTFP1 deletion in cardiomyocytes accelerates, rather than retards, the cardiotoxic effects of doxorubicin (Figure S4F-J), which is consistent with the PCD sensitivity we measured in MEFs and primary cardiomyocytes, we cannot exclude that different cell lines and tissue may respond differently to the loss of MTFP1.

Indeed, MTFP1 ablation significantly impacts mitochondrial respiration in cardiac cells but not in highly glycolytic epithelial cells such as MEFs and human osteosarcoma cells (U2OS) (Figure S2J-Q), in which the bioenergetic efficiency was unaffected.

Efficiently coupled oxidative phosphorylation requires proton gradient formed across the IMM by the mitochondrial ETC, which is then harnessed by ATP synthase to generate ATP from ADP and inorganic phosphate, imported into the matrix by the ADP/ATP translocase (ANT) and phosphate carrier (SLC25A3), respectively. Mitochondrial uncoupling occurs when proton motive force is dissipated by proton leak into the matrix and oxygen consumption is not coupled to ATP generation. Maintenance of constant cellular ATP concentration is critical for cell survival and the function, therefore, uncoupling of the ETC from ATP generation can have deleterious effects in cardiac cells, whose constant energy supply is essential for the beating heart.

While the oxygen consumption rate (OCR) (Figure S2J-Q) and the maximal phosphorylating respiration (Figure 2D, 2H, S2D, S2F), equivalent to state 3, were normal in cells and cardiac mitochondria isolated from cMKO mice, MTFP1 loss significantly reduced the respiratory control ratios (RCR) (Figure 2D, S2D) in cardiac mitochondria energized with complex I or complex II substrates, suggesting a general mechanism of uncoupling of the respiration from the ATP production as a result of increased proton leak trough the IMM.

Proton leak has marked influence on energy metabolism. Enhancement of this process in various tissues can counteract the deleterious effects of nutrient overload via UCP1-dependent and independent pathways (Roesler and Kazak, 2020) and thus may be beneficial in some settings. In the heart, whose bioenergetic efficiency has evolved to maximize ATP output, excessive proton leak has been shown to drive age-related cardiomyocyte and cardiac dysfunction in mice (Chiao et al., 2020; Zhang et al., 2020a). Studies performed by the Rabinovitch lab clearly demonstrated that ANT-dependent proton leak is increased in cardiomyocytes from old, but not young mice and can be rejuvenated by blocking ANT and reducing sensitivity to mPTP opening.

In line with the notion that increased proton leak is maladaptive for the heart, our study shows for the first time that MTFP1 loss in cardiomyocytes reduces the mitochondrial membrane potential as a result of increased proton leak through the IMM (Figure 2D-E, S2D-G) preceding the onset of cardiomyopathy. We provide direct evidence that ANT is the most likely site of proton leak in cardiac mitochondria, as its inhibition with carboxyatractyloside (CATR) suppresses proton leak and re-establishes normal membrane potential and respiratory control ratios in MTFP1-deficient mitochondria (Figure 2H-J). While we have clear evidence of uncoupling via ANT on one hand and increased sensitivity to mPTP opening and mitochondria swelling on the other, future studies are still required to decipher whether these mechanisms are interdependent and whether they must synergize to drive cardiac decline and heart failure. While a number of genetic mouse models of cardiomyopathy targeting mitochondrial genes have been generated over the last 20 years, to the best of our knowledge cMKO mice represent the first in which bioenergetic efficiency is compromised without affecting maximal respiratory capacity (state 3), thus providing a novel model to study the relevance of cardiac mitochondrial uncoupling and its progressive impact on cardiac homeostasis.

In summary our study reveals new and essential roles of MTFP1 in cardiac homeostasis that are distinct from its previously reported impact on mitochondrial fission, the latter of which our data conclusively show is unaffected in vitro and in vivo. Thus, our work now positions MTFP1 as a critical regulator of mitochondrial coupling through ANT in cardiomyocytes and its loss leads to membrane potential dissipation associated to mPTP opening, cell death and progressive DCM that leads to heart failure and middle-aged death. These findings advance our understanding of the mitochondrial defects that can trigger the development of dilated cardiomyopathy and heart failure. We propose that MTFP1 to be a valuable tool for the molecular dissection of mitochondrial uncoupling and mPTP function and thus a promising target to mitigate the pathological events of cardiac and metabolic remodeling in heart disease.

---

***Figure S1. Mtfp1* deletion in cardiomyocytes causes dilated cardiomyopathy and middle-aged death in mice**

**A)** Genotype tissue expression plot (GTEx) of MTFP1 in human tissues.

**B)** Alkaline carbonate extraction assay demonstrating solubility of MTFP1 similar to other multi-pass integral membrane proteins. Immunoblot analysis of soluble (S) and insoluble (pellet, P) fractions of wild type (WT) cardiac mitochondria extracted with Na_2_CO_3_ at the indicated pH. SDHA, SDHB and MT-CO2, ANT1, NDUFA9 were used as markers for soluble and integral membrane proteins, respectively.

**C)** Determination of sub-mitochondrial localization of MTFP1 in cardiac mitochondria by protease K protection assay. TOMM40 and MFN2, were used as OMM markers, CYT C as an IMS marker, and ANT1, ATP5B and SDHA as IMM markers.

**D)** Targeting strategy for conditional inactivation of mouse *Mtfp1*. To allow deletion of *Mtfp1* exons 2 and 3 were flanked in both cases by LoxP sites (blue arrowheads). Flox denotes NeoR cassette containing LoxP targeted locus. LoxP denotes NeoR cassette-deleted targeted locus. Δ denotes deletion induced by Cre-recombinase. FRT sites (green) initially flank NeoR cassette (yellow).

**E)** Kaplan-Meier survival curve of WT (n=11) and cMKO (n=14) female mice. Median lifespan of cMKO mice is 37.5 weeks. Log-rank (Mantel-Cox) test, ***p<0.001.

**F)** M-Mode echocardiography images of WT (top) and cMKO (bottom) female mice at 34 weeks.

**G)** Quantification of left ventricular ejection fraction (% LVEF) of WT (n=7) and cMKO (n=6) female mice at 34 weeks. Data represent mean ± SD. 2-tailed unpaired Student’s t-test, ****p<0.0001.

**H)** Quantification of systolic interventricular septum thickness (IVSs, mm) of WT (n=7) and cMKO (n=6) female mice at 34 weeks. Data represent mean ± SD. 2-tailed unpaired Student’s t-test, *p<0.05.

**I)** Quantification of systolic left ventricular posterior wall thickness (LVPWs, mm) of WT (n=7) and cMKO (n=6) female mice at 34 weeks. Data represent mean ± SD. 2-tailed unpaired Student’s t-test, ***p<0.001.

**J)** Quantification of left ventricle end systolic diameter (LVSD, mm) of WT (n=7) and cMKO (n=6) female mice at 34 weeks. Data represent mean ± SD. 2-tailed unpaired Student’s t-test, ***p<0.001.

**K)** Quantification of left ventricle end diastolic diameter (LVDD, mm) of WT (n=7) and cMKO (n=6) female mice at 34 weeks. Data represent mean ± SD. 2-tailed unpaired Student’s t-test, ***p<0.001.

**L)** Quantification of pulmonary edema. Wet Lung/Body Weight (BW) of WT (n=8) and cMKO (n= 8) at 34 weeks. Data represent mean ± SD; 2-tailed unpaired Student’s t-test, *p<0.05.

**M)** Representative images (left) of WT (top) and cMKO (bottom) hearts isolated from female mice at 34 weeks; (right) quantification of ratio of heart mass to tibia length (mg/mm). Data represent mean ± SD, 2-tailed unpaired Student’s t-test, ***p<0.001.

**N-S)** Isolated primary adult cardiomyocytes from WT (n=51) and cMKO (n=52) mice at 8 weeks were i) field-stimulated at 0.5 Hz, ii) exposed to the β-adrenergic agonist isoproterenol and iii) stimulated at frequency of 5 Hz, iv) and then stepped back to 0.5 Hz washing out (wo) isoproterenol. **N)** Diastolic and systolic sarcomere length (µm) of WT and cMKO; **O**) fractional sarcomere shortening; **P)** Time to 50% or 90% decay of sarcomere shortening in WT or cMKO myocytes, ***p<0.001.

**Q-S)** Isolated primary adult cardiomyocytes from WT (n=29) and cMKO (n=52) mice at 8 weeks were i) field-stimulated at 0.5 Hz, ii) exposed to the β-adrenergic agonist isoproterenol and iii) stimulated at frequency of 5 Hz, iv) and then stepped back to 0.5 Hz washing out (wo) isoproterenol. Intracellular Ca^2+^ concentrations ([Ca^2+^]_c_) were assessed by loading cells with Indo-1 AM and recording calcium transients at 405 nm/485 nm. **Q)** [Ca^2+^]_c_ at diastole or systole; **R)** amplitude of [Ca^2+^]_c_ transients; **S**) Time to 50% or 90% decay (RT) of [Ca^2+^]_c_ in WT and cMKO myocytes, **p<0.01, ****p<0.0001.

***Figure S2. Mtfp1 is required for bioenergetic efficiency in cardiac mitochondria***

**A)** Equal amounts of protein extracted from WT (n=4) and cMKO (n=6) hearts at 18 weeks were separated by SDS–PAGE and immunoblotted with the indicated antibodies.

**B)** Volcano plot of the Label Free Quantification of the cardiac proteome in WT and cMKO mice (18 weeks) listed in Dataset EV1.

**C)** Ratio of NAD(P)H/FAD of the redox states of NAD(P)H/NAD(P)^+^ and FADH_2_/FAD assessed in field-stimulated WT (n=51) and cMKO (n=51) cardiomyocytes isolated from 8-10 weeks old mice (n=3).

**D)** Oxygen consumption rates (left; JO_2_) measured by high-resolution respirometry of cardiac mitochondria isolated from WT (n=5) and cMKO (n=5) mice between 8-10 weeks. Respiration was measured in presence of succinate and rotenone (state 2) followed by the addition of ADP (state 3) and Oligomycin (Omy-state 4). Data represent mean ± SD. Multiple t-test, state 2 ***p<0.001, state 4 *p<0.05. Respiratory control ratios (RCR) of state 3:2 (middle; ADP/Succinate (SUCC)) and state 3:4 (right; ADP/Omy) under complex-II driven respiration. Data represent mean ± SD; multiple t-test, *p<0.05.

**E)** Mitochondrial membrane potential (ΔΨ) measured by quenching of Rhodamine 123 (RH123) fluorescence in cardiac mitochondria from WT (n=8) and cMKO (n=8) mice between 8-10 weeks. ΔΨ was measured in presence of rotenone and succinate (state 2) followed by the addition of ADP (state 3) and Oligomycin (state 4). Data represent mean ± SD; Multiple t-test, *p<0.05, ***p<0.001.

**F)** Oxygen consumption rates (JO_2_) measured by high-resolution respirometry of cardiac mitochondria isolated from WT (n=6) and cMKO (n=6) mice between 8-10 weeks. Respiration was measured in presence of malate (state 2) followed by the addition of ADP (state 3) and palmitoyl-carnitine (PC). Data represent mean ± SD.

**G)** Mitochondrial membrane potential (ΔΨ) measured by quenching of Rhodamine 123 (RH123) fluorescence in cardiac mitochondria isolated from WT (n=6) and cMKO (n=6) mice between 8-10 weeks. ΔΨ sustained by fatty acid oxidation was measured in presence of malate (state 2) followed by the addition of ADP (state 3) and palmitoyl-carnitine (PC). Data represent mean ± SD; Multiple t-test, ***p<0.001.

**H)** PCR genotyping of *Mtfp1* alleles in mice. Deletion of exons 2 and 3 was obtained by crossing *Mtfp1^L^*^oxP/LoxP^ mice with *CMV-Cre* recombinase mice to generate heterozygous knockout *Mtfp1*^+/-^ offspring. Agarose gel of PCR amplicons of genomic DNA. The wild type (387 bp) and the deleted (737 bp) alleles are shown for *Mtfp1^+/+^* and *Mtfp1^-/-^* cells.

**I)** Equal amounts of protein extracted from WT and *MTFP1*^Crispr^ U2OS cells separated by SDS–PAGE and immunoblotted with antibodies against MTFP1 and normalized against Stain Free as a loading control.

**J-M**) Mitochondrial respiration measured in adherent wild type (WT) and *Mtfp1^-/-^* immortalized mouse embryonic fibroblasts (MEFs) cells using the Seahorse Flux Analyzer. **J)** Oxygen consumption rate (OCR) normalized to protein concentration under **K)** Basal **L)** Maximal (FCCP) and **M)** non-phosphorylating (state 4) respiration (Oligomycin treated - Omy). Data are means ± SD of n=4 independent experiments measured on different days. Each point represents the mean of 7-12 technical OCR measurements replicates of each independent experiment.

**N-Q)** Mitochondrial respiration measured in adherent wild type (WT) and *MTFP1^Crispr^* human U2OS osteosarcoma cells using the Seahorse Flux Analyzer. **N)** Oxygen consumption rate (OCR) normalized to protein concentration under **O)** Basal **P)** Maximal (FCCP) and **Q)** non-phosphorylating (state 4) respiration (Oligomycin treated - Omy). Data are means ± SD of 7-12 technical replicates.

**R)** Inhibition of proton leak by MTFP1. Proposed model of the regulation of ANT-dependent proton leak by MTFP1 (red). Proton gradient generated by the electron transport chain (ETC) is harnessed by ATP synthase in WT cells to generate ATP from ADP and Pi. Upon Mtfp1 ablation (*Mtfp1^-/-^*), ANT-mediated proton leak increases thus compromising bioenergetic efficiency.

***Figure S3. MTFP1 is dispensable for mitochondrial fission.***

**A)** Immunoblot of mitochondrial fission and fusion proteins in WT and *Mtfp1^-/-^* MEFs. Equal amounts of protein extracted from WT and *Mtfp1*^-/-^ MEFs were separated by SDS–PAGE and immunoblotted with the indicated antibodies (horizontal line denotes different membranes). VINCULIN was used as loading control.

**B)** Representative confocal images of fragmented mitochondria morphology in MEFs stably expressing FLAG-MTFP1 (mitoYFP, green). EV (empty vector, top), FLAG-MTFP1 (bottom). Scale bar=100 μm.

**C)** Representative immunoblot of mitochondrial fission (red) and fusion (green) proteins in WT and FLAG-MTFP1 stably overexpressing MEFs. Equal amounts of protein were separated by SDS– PAGE and immunoblotted with the indicated antibodies (horizontal line denotes different membranes).

***Figure S4. MTFP1 protects against mitochondrial PTP opening and cell death***

**A)** Mitochondrial permeability transition pore (mPTP) opening. Swelling assay performed on mitochondria purified from WT and *Mtfp1^-/-^* MEFs. Mitochondrial absorbance changes (absorbance 540 nm, % relative to T_0_) are measured every 20 s prior and after addition of a single pulse of CaCl_2_ (arrowhead). Cyclosporin A (CsA) was added prior Ca^2+^ to inhibit mPTP-dependent swelling. Data are means ± SD.

**B)** Representative confocal images of WT and *Mtfp1^-/-^* MEFs subjected to staurosporine (STS) treatment in the presence or absence of the pan-caspase inhibitor q-VD-OPh hydrate (qVD). Cell death was monitored by measuring Propidium Iodide uptake (PI) and imaging cells every hour (h) for 18 h. Scale bar = 100 μm.

**C)** Kinetics of PI uptake was determined by counting the number of PI^+^ positive cells (orange) over total number cells nuclear stained with NucBlue (NB, blue) and expressed as % PI^+^/NucBlue. Data are means ± SD of n=4 independent experiments.

**D)** one-way ANOVA of C) at 12h, ****p<0.0001.

**E)** Cell proliferation curves of WT and *Mtfp1^-/-^* MEFs in glucose (4.5 g/L) containing medium. Number of cells was determined by counting the number of NucBlue (NB) labeled nuclei every 24 h.

**F)** Representative M-Mode echocardiographic images of left ventricles of WT (left) and cMKO (right) mice treated with doxorubicin (Doxo, cumulative 20 mg/kg) at 10 weeks.

**G)** Quantification of left ventricular ejection fraction (% LVEF) of WT (n=5) and cMKO (n=5) after 2 weeks of Doxo administration. Data represent mean ± SD. 2-tailed unpaired Student’s t-test, *p<0.05.

**H)** Quantification of systolic left ventricular diameter (LVSD, mm) of WT (n=5) and cMKO (n=5) mice after Doxo treatment. Data represent mean ± SD. 2-tailed unpaired Student’s t test, *p<0.05.

**I)** Quantification of diastolic left ventricular diameter (LVDD, mm) of WT (n=5) and cMKO (n=5) mice after Doxo treatment. Data represent mean ± SD. 2-tailed unpaired Student’s t test, *p<0.05.

**J)** Quantification of systolic left ventricular posterior wall thickness (LVPWs, mm) of WT (n=5) and cMKO (n=5) mice after Doxo treatment. Data represent mean ± SD. 2-tailed unpaired Student’s t test, *p<0.05.

***Figure S5. Generation of FLAG-MTFP1 mouse model to define the cardiac interactome***

**A)** For inducible overexpression of mCherry-P2A-Flag-Mtp18, two mouse models were generated from one construct. The first (upper pane) allowing inducible expression from the CAG-promoter, the second (lower panel) from the mouse Rosa26 promoter. To do so, the construct in the upper panel is generated and recombined into the mouse Rosa26 locus. The genuine Rosa26 promoter is further upstream, and its transcript is spliced into the splice acceptor of exon 2 (“s.acc.”, purple). In the native state, transcription is aborted at the insulator site (H19, black), and instead started at the CAG promoter (pCAG, yellow arrow). The loxP-flanked (lox, blue arrow heads) neo–STOP cassette for conditional activation has a stop site that is a large region of SV40 intron plus late polyA (neo, green arrow; STOP, blue box). The cDNA is inserted downstream of the neo/stop cassette. Both, Rosa26-driven and CAG-driven expression, is polyadenylated at a bovine growth hormone polyA site (bGH pA, small black box). For exchange of the promoters, the CAG-cassette is flanked by FRT-sites. This cassette will be excised by Flp-mediated deletion in vivo, leaving mCherry-P2A-Flag-Mtp18 expression under control of the Rosa26 promoter. Upon Cre mediated deletion, the mCherry-P2A-Flag-Mtp18 cDNA is expressed from the CAG promoter (upper panel) or the intrinsic Rosa26 promoter (lower panel).

**B)** Representative immunoblot of the expression of Flag-MTFP1 compared to endogenous MTFP1 levels in KI mice.

**C)** Quantification of left ventricular ejection fraction (% LVEF) of WT (n=4) and KI (n=3) mice at 20 weeks. Data represent mean ± SD.

**D)** Volcano plot of the FLAG-MTFP1 interactome analyzed by mass spectrometry. Mitochondrial proteins exclusively present in FLAG-MTFP1 eluates or significantly enriched greater than two-fold, listed in Dataset EV3.

**E)** Functional classification of 60 mitochondrial proteins identified in Co-IP eluates in C) (Dataset EV3).

## Material and Methods

### Animals

Animals were handled according to the European legislation for animal welfare (<u>Directive 2010/63/EU</u>). All animal protocols were reviewed and approved by the local and national authorities. Mice were housed within a specific pathogen free facility and maintained under standard housing conditions of a 14-10h light-dark cycle, 50-70% humidity, 19-21°C with free access to food and water in cages enriched with bedding material and gnawing sticks. *Mtfp1* conditional mice (*Mtfp1*^LoxP/LoxP^) were generated by PolyGene AG (Switzerland) on a C57Bl6/N background. Cardiomyocyte specific *Mtfp1* KO mice (cMKO; *Myh6-Cre^tg/+^Mtfp1^LoxP/LoxP^*) were generated by crossing *Mtfp1* conditional mice (*Mtfp1^LoxP/LoxP^*) with transgenic (Tg) mice expressing the Cre recombinase under the control of the cardiac alpha myosin heavy chain 6 promoter (*Myh6-Cre*)(Agah et al., 1997). Littermates that were homozygous for the conditional allele and negative for *Myh6-Cre* were used as controls (WT; *Myh6-Cre^+/+^Mtfp1^LoxP/LoxP^*). Heterozygous whole-body *Mtfp1* KO mice (*Mtfp1^+/-^*) were generated by crossing *Mtfp1* conditional mice (*Mtfp1^LoxP/LoxP^*) with transgenic (Tg) mice expressing the Cre recombinase under the control of the CMV promoter (*CMV-Cre*)(Schwenk et al., 1995) and backcrossing to C57Bl6/N wild type (WT) mice.

Cardiomyocyte specific FLAG-MTFP1 Knock-In (KI) mice (*Myh6-Cre^tg/+^Mtfp1^+/+^, CAG^tg+/^)* were generated by crossing an inducible mouse model for mCherry-P2A-FLAG-MTFP1 generated by PolyGene AG (Switzerland) on a C57Bl6/N background expression under the CAG promoter with mice expressing the Cre recombinase under the control of the cardiac alpha-(α) myosin heavy chain 6 promoter (*Myh6-Cre*).

### Cell Lines

WT (*Mtfp1^+/+^*) and knockout (KO) *Mtfp1^-/-^* embryos were isolated at E13.5 following F1 heterozygous intercrosses of *Mtfp1^+/-^* whole-body KO mice. Immortalization of WT (*Mtfp1^+/+^*) and *Mtfp1^-/-^* primary mouse embryonic fibroblasts (MEFs) was performed as previously described(Cretin et al., 2021, 2021) using a plasmid encoding SV40 large T antigen. MEFs cells were maintained in Dulbecco’s modified Eagle’s medium (DMEM + GlutaMAX, 4.5 g/L D-Glucose, pyruvate) supplemented with 5% FBS and 1% penicillin/streptomycin (P/S, 50 µg/ml) in a 5% CO_2_ atmosphere at 37°C.

Genetic disruption of *Ppif* in WT and *Mtfp1^-/-^* MEFs was performed via CRISPR-Cas9 gene editing targeting Exon 1 of *Ppif* (sgRNA: forward: aaacCCGGGAACCCGCTCGTGTAC and reverse: CACCGTACACGAGCGGGTTCCCGG. sgDNA oligonucleotides were annealed and cloned into pSpCas9(BB)-2A-GFP PX458 to generate pTW363 (pSpCas9(BB)-2A-GFP PPIFsgDNA). WT (*Mtfp1^+/+^*) and *Mtfp1^-/-^* immortalized MEFs were transfected with 2.5 μg of plasmid pTW363 using Lipofectamine 2000 (Life Technologies). After 48h incubation, cells were individually isolated in 96 well plates by FACS. Clones were then expanded and validated by Sanger sequencing and western blotting. Both single Ppif KO MEFs (*Ppif^-/-^*) and double KO *Mtfp1^-/-^Ppif^-/-^* cells carry the same homozygous c.126delG insertion that is predicted to yield a truncated polypeptide product at amino acid position 65 (1-64 protein) p.Val65*. Immunoblot analysis was used to confirm the absence of PPIF protein.

MEFs expressing mitochondrially targeted YFP (mitoYFP) were generated from Gt(ROSA26)Sor^mitoYFP/+^ embryos on a C57Bl6/N genetic background at E13.5 and immortalized using a plasmid encoding SV40 large T antigen as previously described(Cretin et al., 2021). MEFs stably expressing FLAG-MTFP1 were generated by lentiviral transduction with pTW142 (pLVX-EF1α-MTFP1) containing a puromycin resistant marker. The empty vector (EV) pLVX-EF1α (pTW122) was used to generate control cells.

Human U2OS osteosarcoma cells were depleted for MTFP1 by CRISPR-Cas9 gene editing. The single-guide RNAs (sgRNAs) were designed using the CRISPR-Cas9 design tool (benchling.com) to target Exon 1 of *MTFP1*. sgDNA oligonucleotides (forward: 5’-caccgGCGCAGAGCGCGATCTCTAC-3’ and reverse: 5’-aaacGTAGAGATCGCGCTCTGCGCc-3’) were annealed and cloned into the BbsI digested pSpCas9(BB)-2A-GFP vector (SpCas9(BB)-2A-GFP (PX458) which was a gift from Feng Zhang (Addgene plasmid # 48138). U2OS cells were transfected in 6-well dishes with 2ug of pSpCas9(BB)-2A-GFP plasmid containing the respective sgRNA using Lipofectamine 2000 (Life Technologies, 11668027). After 24h incubation, GFP positive cells were individually isolated by fluorescence-activated cell sorting. Clones were expanded and were validated by western blotting and DNA sequencing.

### SDS-PAGE immunoblot analysis

Immunoblot analysis was used to assess steady-state protein levels in cardiac tissue and cell lysates. For tissue lysates, mice were sacrificed by cervical dislocation, the chests were opened, and the hearts were excised, weighed, flash frozen in liquid nitrogen, and stored at −80°C until use. The LV tissue or MEFs cellular pellet was homogenized in cold RIPA buffer [1 mg/ 20 µL, 1% Triton X-100, 1% sodium deoxycholate, 0.1% SDS, 150 mM NaCl, 50 mM Tris·HCl (pH 7.8), 1 mM EDTA, and 1 mM EGTA] in presence of protease and phosphatase inhibitors and kept on ice for 30 minutes. The homogenate was then centrifuged for 15 min at 16,000g, 4°C. The protein concentration was determined by Bradford assay (Bio-Rad) using a BSA standard curve. The protein absorbance was measured at 595 nm by using a microplate reader Infinite M2000 (Tecan). Equal amounts of protein were reconstituted in 4x Laemmli Sample Buffer [355 mM, 2-mercaptoethanol, 62.5 mM Tris-HCl pH 6.8, 10% (v/v) glycerol, 1%(w/v) SDS, 0.005% (v/v) Bromophenol Blue] and heated at 95°C for 5 min. Samples (10 µg) were resolved on 4-20% polyacrylamide gels (Mini Protean TGX Stain-Free gels, BioRad) and transferred to nitrocellulose membrane with Trans-Blot® Turbo™ Transfer system (Bio-Rad). Equal protein amount across membrane lanes were checked by Ponceau S staining or Stain-free detection. Membranes were blocked for at least 1h with 5% (w/v) semi-skimmed dry milk dissolved in Tris-buffered saline Tween 0.1% (TBST), incubated overnight at 4°C with primary antibodies dissolved 1:1,000 in 2% (w/v) Bovine Serum Albumin (BSA), 0.1% TBST. The next day membranes were incubated in secondary antibodies conjugated to horseradish peroxidase (HRP) at room temperature for 2h (diluted 1:10,000 in BSA 2% TBST 0.1%). Finally, membranes were incubated in Clarity™ Western ECL Substrate (Bio-Rad) for 2 min and luminescence was detected using the ChemiDoc® Gel Imaging System. Densitometric analysis of the immunoblots was performed using Image Lab Software (Bio-Rad).

### Mitochondrial isolation

Isolated cardiac mitochondria were freshly isolated as previously described (Mourier et al., 2014) with some modifications. Briefly, the heart was washed in ice-cold PBS solution. Ventricles were separated from atria and non-myocardial tissue, cut in small pieces, and then transferred to an ice-cold 2-ml homogenizer (Teflon pestle) and manually homogenized in IB buffer (sucrose 275 mM, Tris 20 mM, EGTA-KOH 1 mM, pH 7.2) containing Trypsin-EDTA (0.05%). Trypsin activity was then inhibited by adding to the homogenate bovine serum albumin (BSA) fatty acid free (0.25 mg/mL) and protease inhibitor cocktail (PIC, Roche).

Mitochondria isolation from MEFs was performed starting from 10 x 150 mm dishes at 100% confluence. Cells were collected into 10 mL of IB Buffer containing BSA fatty acid free (0.25 mg/mL) and PIC and homogenized with 30 stokes of the plunger at 1500 rpm on ice.

Cardiac and/or cellular homogenates were then centrifuged at low speed (1000 g, 10 min, 4 °C) to discard nuclei and debris, and further centrifuged (3200 g, 15 min, 4°C) to obtain the crude mitochondrial pellet and the cytosolic fraction. The crude mitochondrial pellet was finally resuspended in IB buffer containing BSA and PIC and protein concentration was determined by using the Bradford assay.

### Protease Protection Assay

Crude mitochondria isolated from WT mouse hearts was subjected to protease protection assay as previously described (Anand et al., 2014). 50 µg of crude mitochondria were resuspended into the following buffers without protease inhibitors: 1) Mitochondrial isolation IB buffer 2) Mitochondrial isolation buffer with Proteinase K (100 µg/mL) 3) Swelling buffer (EDTA 1 mM, HEPES 40 mM) 4) Swelling buffer + Proteinase K (100 µg/mL) 5) Swelling buffer + Proteinase K (100 µg/ml) + Triton X-100 0.5% and incubated at 37°C for 30 minutes at 750 rpm. Mitochondrial proteins were then precipitated in trichloric acid buffer (14% TCA, 40 mM HEPES, 0.02% Triton X-100) for 3 hours at −20°C and centrifugated for 20 min. Pellets were washed twice with ice-cold acetone and air-dried prior to re-suspension in 1x Laemilli sample buffer (Bio-Rad) for SDS-PAGE and immunoblot analysis.

### Alkaline Carbonate Extraction

Alkaline carbonate extraction of membrane proteins was performed as previously described (Anand et al., 2014). Crude mitochondria isolated from WT mouse hearts were resuspended and incubated for 30 min on ice with 0.1 M Na_2_CO_3_ at the following pH: 12.5, 11.5, 10.5, or 9.5. The suspensions were ultra-centrifuged at 4°C for 30 min at 90,000 g in Beckman polycarbonate tubes in a TLA 110 rotor. Supernatants and pellets were then incubated in trichloric acid buffer (14% TCA, 40 mM HEPES, 0.02% Triton X-100) for 15 min on ice, followed by centrifugation for 20 min at 28,000 g at 4°C. The samples were washed 3x with 100% acetone, subsequently dried for 30 min at RT. The dried pellet was then resuspended with 1x Laemilli sample buffer (Bio-Rad) for SDS-PAGE and western blot analysis.

### siRNA transfection

Silencing of the indicated genes was performed using forward transfection: 20 nM of the indicated SmartPool siRNAs listed in Dataset EV4 were mixed with Lipofectamine RNAiMax (Invitrogen), added on top of cells seeded in 6-well dishes (300,000 cells/well) and incubated at 37°C in a CO_2_ incubator. For live imaging experiments, cells were seeded in 96 well plates (Cell Carrier Ultra, Perkin Elmer) 48h after transfection to perform live imaging after 72h post-transfection.

### Cardiac RNA sequencing and RT-qPCR

WT and cMKO mice aged 8 and 18 weeks were sacrificed by cervical dislocation and hearts were quickly excised and rinsed with cold sterile PBS. Left ventricle posterior walls were collected and snap frozen in liquid nitrogen and stored at −80°C until use. Total RNA was extracted by using TRIzol (Invitrogen, NY, USA) according to standard procedures. The Trizol/chloroform mixture was centrifuged at 12000g for 15 min at 4°C, the supernatant corresponding to the aqueous phase was transferred to a new tube, mixed with ethanol 70%, and then applied to an RNA mini-column extraction kit (NucleoSpin RNA kit, MACHEREY-NAGEL) according to the manufacturer’s recommendations. After DNA digestion, total RNA was eluted with RNase-free water and stored in liquid nitrogen. RNA was quantified by using the Bioanalyzer 2100 (Agilent) and only RNA samples (≥ 50 ng/µL) with an RNA integrity value (RIN) ≥ 6.8, OD^260^/^280^ ≥ 2, OD^260^/^230^ ≥ 2 were considered for the transcriptome profiling. For RNA sequencing, libraries were built using a TruSeq Stranded mRNA library Preparation Kit (Illumina, USA) following the manufacturer’s protocol. Two runs of RNA sequencing were performed for each library on an Illumina NextSeq 500 platform using single-end 75bp. The RNA-seq analysis was performed with Sequana 0.8.5(Cokelaer et al., 2017) using an RNA-seq pipeline 0.9.13 (https://github.com/sequana/sequana_rnaseq) built on top of Snakemake 5.8.1(Köster and Rahmann, 2012). Reads were trimmed from adapters using Cutadapt 2.10(Martin, 2011) then mapped to the mouse reference genome GRCm38 using STAR 2.7.3a(Dobin et al., 2013). FeatureCounts 2.0.0 was used to produce the count matrix, assigning reads to features using annotation from Ensembl GRCm38_92 with strand-specificity information(Liao et al., 2014). Quality control statistics were summarized using MultiQC 1.8(Ewels et al., 2016). Statistical analysis on the count matrix was performed to identify differentially expressed genes (DEGs), comparing WT and cMKO. Clustering of transcriptomic profiles were assessed using a Principal Component Analysis (PCA). Differential expression testing was conducted using DESeq2 library 1.24.0 scripts based on SARTools 1.7.0(Varet et al., 2016) indicating the significance (Benjamini-Hochberg adjusted p-values, false discovery rate FDR < 0.05) and the effect size (fold-change) for each comparison.

For RT-qPCR, 1 µg of total RNA was converted into cDNA using the iScript Reverse Transcription Supermix (Bio-Rad). RT-qPCR was performed using the CFX384 Touch Real-Time PCR Detection System (Bio-Rad) and SYBR® Green Master Mix (Bio-Rad) using the primers listed in Dataset EV1. Gapdh was amplified as internal standard. Data were analyzed according to the 2−ΔΔCT method(Livak and Schmittgen, 2001).

### Proteomics

Heart extract proteins were extracted and denatured in RIPA buffer. Samples were sonicated using a Vibracell 75186 and a miniprobe 2 mm (Amp 80% // Pulse 10 off 0.8, 3 cycles) and further centrifuged. Protein assay was performed on supernatant (Pierce 660 nm, according to manufacturer instructions) and 100 µg of each extract was delipidated and cleaned using a Chloroform / Methanol / Water precipitation method. Briefly, 4 volume of ice-cold methanol were added to the sample and vortex, 2 volume of ice-cold chloroform were added and vortex and 3 volume of ice-cold water was added and vortex. Samples were centrifuged 5 min at 5000 g. The upper layer was removed and proteins at the interface were kept. Tubes were filled with ice-cold methanol and centrifuged at max speed for 5 min. Resulting protein pellet was air-dried and then dissolved in 130 µl of 100mM NaOH before adding 170 µl of Tris 50 mM pH 8.0, tris (2-carboxyethyl)phosphine (TCEP) 5 mM and chloroacetamide (CAA) 20 mM. The mixture was heated 5 min at 95 °C and then cooled on ice. Endoprotease LysC (1µg) was use for a 8h digestion step at (37° C) followed with a trypsin digestion (1 µg) at 37°C for 4 h. Digestion was stopped adding 0.1% final of trifluoroacetic acid (TFA). Resulting peptides were desalted using a C18 stage tips strategy (Elution at 80% Acetonitrile (ACN) on Empore C18 discs stacked in a P200 tips) and 30 µg of peptides were further fractionated in 4 fractions using poly(styrenedivinylbenzene) reverse phase sulfonate (SDB-RPS) stage-tips method as previously described(Kulak et al., 2014; Rappsilber et al., 2007). Four serial elutions were applied as following: elution 1 (80 mM Ammonium formate (AmF), 20% (v/v) ACN, 0.5% (v/v) formic acid (FA)), elution 2 (110 mM AmF, 35% (v/v) ACN, 0.5% (v/v) FA), elution 3 (150 mM AmmF, 50% (v/v) ACN, 0.5% (v/v) FA) and elution 4 (80% (v/v) ACN, 5 % (v/v) ammonium hydroxide). All fractions were dried and resuspended in 0.1% FA before injection.

#### NanoLC-MS/MS

LC-MS/MS analysis of digested peptides was performed on an Orbitrap Q Exactive Plus mass spectrometer (Thermo Fisher Scientific, Bremen) coupled to an EASY-nLC 1200 (Thermo Fisher Scientific). A home-made column was used for peptide separation (C_18_ 50 cm capillary column picotip silica emitter tip (75 μm diameter filled with 1.9 μm Reprosil-Pur Basic C_18_-HD resin, (Dr. Maisch GmbH, Ammerbuch-Entringen, Germany)). It was equilibrated and peptide were loaded in solvent A (0.1 % FA) at 900 bars. Peptides were separated at 250 nl.min^-1^. Peptides were eluted using a gradient of solvent B (ACN, 0.1 % FA) from 3% to 7% in 8 min, 7% to 23% in 95 min, 23% to 45% in 45 min (total length of the chromatographic run was 170 min including high ACN level step and column regeneration). Mass spectra were acquired in data-dependent acquisition mode with the XCalibur 2.2 software (Thermo Fisher Scientific, Bremen) with automatic switching between MS and MS/MS scans using a top 12 method. MS spectra were acquired at a resolution of 35000 (at *m/z* 400) with a target value of 3 × 10^6^ ions. The scan range was limited from 300 to 1700 *m/z*. Peptide fragmentation was performed using higher-energy collision dissociation (HCD) with the energy set at 27 NCE. Intensity threshold for ions selection was set at 1 × 10^6^ ions with charge exclusion of z = 1 and z > 7. The MS/MS spectra were acquired at a resolution of 17500 (at *m/z* 400). Isolation window was set at 1.6 Th. Dynamic exclusion was employed within 45 s.

#### Data Processing

Data were searched using MaxQuant (version 1.5.3.8) using the Andromeda search engine(Tyanova et al., 2016) against a reference proteome of Mus musculus (53449 entries, downloaded from Uniprot the 24^th of^ July 2018). The following search parameters were applied: carbamidomethylation of cysteines was set as a fixed modification, oxidation of methionine and protein N-terminal acetylation were set as variable modifications. The mass tolerances in MS and MS/MS were set to 5 ppm and 20 ppm respectively. Maximum peptide charge was set to 7 and 5 amino acids were required as minimum peptide length. A false discovery rate of 1% was set up for both protein and peptide levels. All 4 fractions per sample were gathered and the iBAQ intensity was used to estimate the protein abundance within a sample(Schwanhäusser et al., 2011). The match between runs features was allowed for biological replicate only.

For this large-scale proteome analysis part, the mass spectrometry data have been deposited at the ProteomeXchange Consortium (http://www.proteomexchange.org) via the PRIDE partner repository(Perez-Riverol et al., 2019; Vizcaíno et al., 2014) with the dataset identifier PXD028516

### mtDNA content quantification

Genomic DNA was extracted using the NucleoSpin Tissue (MACHEREY-NAGEL) and quantified with NanoQuant Plate™ (Infinite M200, TECAN). RT-qPCR was performed using the Real-Time PCR Detection System (Applied Biosystems StepOnePlus), 20 ng of total DNA and the SYBR® Green Master Mix (Bio-Rad). b-*Actin* was amplified as an internal, nuclear gene standard as previously described(Cretin et al., 2021). PCR primer sequences are listed in Dataset EV4. Data were analyzed according to the 2^−ΔΔCT method(Livak and Schmittgen, 2001).

### Mitochondrial imaging and quantification in MEFs

MEFs cells were seeded on 96-well or 384-well CellCarrier Ultra imaging plates (Perkin Elmer) 24h before imaging. Nuclei were stained with NucBlue™ Live ReadyProbes™ Reagent (ThermoFisher Scientific) at 1 drop per 10 mL of media. Fluorescent labeling of mitochondria was achieved using MitoTracker DeepRed (MTDR) at 100 nM for 30 min at 37°C, 5% CO_2_. Cells were then washed with regular medium and images were acquired using the Operetta CLS or Opera Phenix High-Content Analysis systems (Perkin Elmer), with 20x Water/1.0 NA, 40x Air/0.6 NA, 40x Water/1.1 NA or 63x Water/1.15 NA. MTDR (615-645 nm) and NucBlue (355-385 nm) were excited the appropriate LEDs (Operetta CLS) or lasers (Opera Phenix). Automatic single-cell classification of non-training samples (i.e. unknowns) was carried out by the supervised machine-learning (ML) module using Harmony Analysis Software (PerkinElmer) as previously described(Cretin et al., 2021).

### Analysis of oxygen consumption

Oxygen consumption of intact cells or crude mitochondria was measured with the XFe96 Analyzer (Seahorse Biosciences) and High Resolution Respirometry (O2k-Fluorespirometer, Oroboros, AT), respectively. For Seahorse experiments, cells (experimentally optimized density of 20,000 cells/well for MEFs and U2OS cells) were seeded onto 96-well XFe96 cell culture plates. On the following day, cells were washed and incubated with Seahorse XF Base Medium completed with 1 mM Pyruvate, 2 mM Glutamine and 10 mM Glucose. Cells were washed with the Seahorse XF Base Medium and incubated for 45 min in a 37°C non-CO_2_ incubator before starting the assay. Following basal respiration, cells were treated sequentially with: oligomycin 1 µM, CCCP 2 µM and antimycin A plus rotenone (1 µM each) (Sigma). Measurements were taken over 2-min intervals, proceeded by a 1-min mixing and a 30s incubation. Three measurements were taken for the resting OCR, three for the non-phosphorylating OCR, three for the maximal OCR and three for the extramitochondrial OCR. After measurement, the XFe96 plate was washed with Phosphate-Buffered Saline (PBS) and protein was extracted with RIPA for 10 min at RT. Protein quantity in each well was then quantified by Bicinchoninic acid assay (BCA) by measuring absorbance at 562 nm and used to normalize OCR measurements as previously described(Cretin et al., 2021).

For O2k respirometry, isolated cardiac mitochondria were freshly isolated from adult WT and cMKO mice (8, 18 or 34 weeks) as described above. Simultaneous measurement of mitochondrial respiration and membrane potential (Δ*ψ*) was assessed by O2K-Fluorometry using a O2K-Fluorescence LED2-Module operated through the amperometric channel of the O2K. Briefly, 25 or 50 µg of cardiac mitochondria were resuspended in Mir05 buffer [MgCl_2_-6H_2_O 3 mM, Lactobionic Acid 60 mM, Taurine 20 mM, KH_2_PO_4_ 10 mM, Hepes-KOH 20 mM, Sucrose 110 mM, EGTA-KOH 0.5 mM, BSA (1g/L)] in presence of Rhodamine 123 (RH-123) (0.66 µM). Maximal mitochondrial respiration capacity (OXPHOS) was determined under consecutive administrations of PGM [pyruvate 10 mM, glutamate 5 mM, malate 5 mM, state 2] or malate (2 mM) and palmitoyl-carnitine (10 µM) in presence of ADP (1 mM, state 3) to assess complex I-driven respiration; rotenone (0.5 µM) and succinate (10 mM, state 2) for the measurement of complex II-driven respiration; antimycin A (2.5 µM), ascorbate (2 mM), *N*,*N*,*N*′,*N*′-tetramethyl-*p*-phenylenediamine (TMPD, 0.5 mM) and carbonyl cyanide m-chlorophenyl hydrazone (CCCP, 2 µM) for the determination of the complex IV-driven respiration. The addition of oligomycin (Omy, 25 nM) or carboxyatractyloside (CATR, 1 µM) was used to determine the non-phosphorylating respiration (state 4). Cytochrome *c* (2 µM) – mediated O_2_ flux was used to evaluate the integrity of the OMM following the mitochondria isolation. Respiratory control ratios (RCR) of state 3: state 4 were used to evaluate the integrity of the IMM. RH-123 fluorescence quenching (Δ fluorescence) in energized mitochondria was used as a direct measurement of Δ*ψ*.

### Echocardiography

Transthoracic echocardiographic acquisitions were performed by using a Vevo 3100 Imaging System coupled to a 25-55 MHz linear-frequency transducer (MX550D, FUJIFILM VisualSonics). Randomized WT and cMKO mice [10 to 34 weeks of age (male), 34 weeks (female) were anesthetized and maintained with 2% isofluorane in oxygen, placed in a supine position on a 37°C warmed pad. Limb electrodes and a rectal probe were used to monitor define (ECG) and body temperature. Before echocardiography and the addition of the pre-warmed ultrasound gel, thorax fur was removed using hair-removal cream. To assess the overall left ventricle (LV) size and LV function, B-and M-Mode images were acquired in parasternal long axis view (PLAX) between 400-500 bpm. The systolic and diastolic LV dimensions [interventricular septum thickness (IVS; mm), LV diameter (LVD; mm), LV posterior wall thickness (LVPW; mm)] and cardiac output [ejection fraction (% EF)] were determined by acquiring and analyzing at least 3 independent cardiac cycles within at least 3 M-Mode images.

### Doxorubicin Treatment

WT and cMKO mice aged 8 weeks received a cumulative dose of 20 mg/kg of doxorubicin (Doxo, 2 intra-peritoneal injections of 10 mg/kg/saline, given at 3-day intervals; control mice received saline injections). Transthoracic echocardiography was performed at day 14 post treatment. Body weight was recorded every 3 days and a loss higher than 20% was considered as endpoint of the study.

### Determination of serum levels of cardiac troponin I and cardiac MLC1

A mouse ELISA kit was used to compare serum levels of cardiac troponin I (cTnI, Life Diagnostics) and cardiac myosin light chain 1 (MLC1, Life Diagnostic) in WT and cMKO male mice aged 18 and 34 weeks. Blood was collected via submandibular vein puncture from non-anesthetized mice, left for 30 min at RT and then centrifuged at 5,000 g at 4°C for 10 min then snap-frozen into liquid nitrogen. Serum was stored at −80°C until next use. The assays to determine cTnI and MLC1 levels were performed following the exact manufacturer’s instructions.

### Histology

WT and cMKO mice aged 34 weeks were sacrificed by cervical dislocation and the hearts excised post-mortem. The whole hearts were then fixed with Formol 4 % (VWR chemicals) overnight and then fully dehydrated by a series of ethanol gradients. Tissues were then paraffin-embedded and sectioned in a short view at a thickness of 4 μm on a microtome. The sections were de-paraffinized in xylene and rehydrated followed by hematoxylin and eosin (H&E), Masson’s trichrome or Picrosirius Red staining according to the standard protocols.

### Transmission Electron Microscopy

Transmission electron microscopy was performed on cardiac tissue from WT and cMKO mice aged 8-10 weeks. Small pieces (1×1×1 mm) from the left ventricle posterior wall were fixed in a 37°C prewarmed mix of PHEM 1x buffer (60 mM PIPES, 25 mM HEPES, 10 mM EGTA, 2 mM MgCl_2_; pH 7.3), 2.5% glutaraldehyde and 2% PFA for 30 min, followed by an overnight fixation at 4°C. Specimens were then rinsed 3 times with PHEM 2X buffer. Samples were incubated with 1 % osmium tetroxide (Merck) and 1.5 % ferrocyanide (Sigma Aldrich) in 0.1M PHEM. After dehydration by a graded series of ethanol, samples were gradually infiltrated at RT with epoxy resin and after heat polymerization, sections with a thickness of 70 nm were cut with a Leica UCT microtome and collected on carbon, formvar coated copper grids. Sections were contrasted with 4% aqueous uranylacetate and Reynold’s lead citrate. Generation of ultra-large electron microscopy montages at a magnification of 14500x (pixel size = 6.194 nm, bin 1, US4000 Ultrascan camera) were acquired using a TECNAI F20 Transmission Electron Microscope (FEI) with a field emission gun (FEG) as an electron source, operated at 200kV. The SerialEM software(Mastronarde and Held, 2017; Schorb et al., 2019) was used for multi-scale mapping as previously described(Cretin et al., 2021). The ‘Align Serial Sections/Blend Montages’ interface of IMOD16 was used for blending the stacks of micrographs collected to single large images. Quantification of mitochondrial cross-sectional area was determined by tracing the perimeter of individual mitochondria with the use of 3dmod software.

### Primary adult cardiomyocytes isolation

Primary adult ventricular cardiomyocytes (CMs) were isolated from WT (*Myh6-Cre^+/+^*, *Mtfp1*^LoxP/LoxP^) and *Mtfp1* KO mice (cMKO, *Myh6-Cre^Tg/+^*, *Mtfp1*^LoxP/LoxP^) mice aged 8-10 weeks to perform live-cell imaging experiments by using the simplified Langendorff-free method as previously published (Ackers-Johnson et al., 2016) with some modifications. Briefly, mice were weighed, injected intraperitoneally with ketamine/xylazine (ketamine: 3 x 80 mg/kg/bw; xylazine 3 x 10 mg/kg/bw). Once the thorax was opened, the descending aorta was cut and the heart exposed and flushed into the right ventricle with EDTA buffer pH 7.8 (NaCl 130 mM, KCl 5 mM, NaH_2_PO_4_ 0.5 mM, HEPES 10 mM, Glucose 10 mM, 2,3-butanedione monoxime (BDM) 10 mM, Taurine 10 mM, EDTA 5 mM). The heart was then excised, transferred into a dish containing EDTA buffer, where the ascending aorta was clamped. Tissue digestion was performed by sequential injection of EDTA buffer, perfusion buffer (NaCl 130 mM, KCl 5 mM, NaH_2_PO_4_ 0.5 mM, HEPES 10 mM, Glucose 10 mM, BDM 10 mM, Taurine 10 mM, MgCl2 1 mM, pH 7.8) and collagenase buffer (Collagenase II 0.5 mg/mL, Collagenase 4 0.5 mg/mL, Protease XIV 0.05 mg/mL) into the left ventricle (LV) through the apex. To control the flow of the digestion buffer the LV was perfused via a flexible linker using an automated infusion pump (Graseby 3200). After digestion, ventricles were then separated from the atria and ultimately dissociated by gentle pipetting. The digestion process was then inhibited by the addition of the stop buffer (perfusion buffer containing FBS 10%). Cellular suspension was filtered using a 100 μm pore size strainer to remove undigested debris. CMs were then collected by gravity settling (15 min) while resuspended sequentially in 3 calcium reintroduction buffers to gradually restore calcium levels (0.34 mM, 0.68 mM, and 1.02 mM Ca^2+^). CMs yields were quantified using a hemocytometer. CMs were resuspended in prewarmed culture media (M199 Medium, BSA 0.1%, Insulin-Transferrin-Selenium (ITS) 1x, BDM 10 mmol/l, CD lipid 1x, P/S 1x), and plated into laminin (10 μg/mL) precoated Cell Carrier Ultra-96 well (Perkin Elmer) plates, in a humidified tissue culture incubator (37°C, 5% CO_2_) for at least 1 hour before proceeding to live imaging.

### Cell death assay

CMs isolated from WT and cMKO mice (8-10 weeks) labelled with NucBlue™ Live ReadyProbes™ Reagent (ThermoFisher Scientific) and Tetramethylrhodamine Ethyl Ester Perchlorate (TMRE; 50 nM) for 20 min at 37°C, 5% CO_2_. Supervised machine learning (Harmony 4.9, Perkin Elmer) using TMRE intensity, cell geometry (rod-shape, area, length) and nuclei number as read-outs enabled the on-the-fly detection, imaging, and quantification of healthy CMs from dead and dying CMs and non-myocytes. Live-cell imaging was performed in the presence or absence of a cell death stimuli: H_2_O_2_ (25 µM), or doxorubicin (Doxo, 60 µM,) or carbonyl cyanide m-chlorophenyl hydrazine (CCCP, 10 µM). Image acquisition was performed using the Operetta CLS High-Content Analysis system (Perkin Elmer) with a 5x air objective.

WT, *Mtfp1*^-/-^, *Ppif*^-/-^ and *Mtfp1*^-/-^*Ppif*^-/-^ MEFs were plated in Cell Carrier Ultra-96 well (Perkin Elmer) and incubated 24h with regular media. Cells were then stained with NucBlue™ Live ReadyProbes™ to visualize nuclei. CellEvent Caspase 3/7 Green (CE) was added (1 drop per 1 mL of media) to visualize the activation of caspases 3/7 in cells undergoing to apoptosis. Propidium Iodide (PI, 1:500) was added to visualize dead cells. Cell death was induced with actinomycin D (1.5 µM) alone, or in association with ABT-737 (10 µM), an inhibitor of the Bcl-2 family proteins, or doxorubicin hydrochloride (DOXO, 1.5 µM), or staurosporine (1 µM) or hydrogen peroxide H_2_O_2_ (500 µM). The pan-caspase inhibitor Q-VD-OPh (qVD, 20 µM) and cyclophilin D inhibitor cyclosporin A (CsA, 2 µM), were added where indicated. Image acquisition was performed using the Operetta CLS High-Content Analysis system (Perkin Elmer) with a 20x air objective. CE or PI positive nuclei number were automatically counted by using Harmony 4.9 software.

### Field-stimulated cardiomyocytes

Adult ventricular cardiomyocytes (CMs) from WT and cMKO mice (8-10 weeks) were isolated with a Langendorff system as previously described (Chen et al., 2012; Nickel et al., 2015) and paced by electrical field stimulation at 37°C using a customized IonOptix system. CMs were exposed to a protocol that simulates a physiological workload increase by first pacing cells at 0.5 Hz in Normal Tyrodés solution pH 7.4 [NT; NaCl 130 mM, KCl 5 mM, MgCl_2_ 1 mM, CaCl_2_ 1 mM, Na-HEPES 10 mM, glucose 10 mM, sodium pyruvate 2 mM and ascorbic acid 0.3 mM]. The β-adrenergic receptor agonist isoproterenol (30 nM) was then washed in for 1 min and then the stimulation rate was increased to 5 Hz for 3 min. After this time, stimulation rate was set back to the initial 0.5 Hz, and isoproterenol washed out. During the measurements or sarcomere length and contraction, the system detection was combined to recordings of the autofluorescence of NAD(P)H/NAD(P)^+^ and FADH_2_/FAD by alternately exciting cells at wavelengths (λ_exc_) of 340 and 485 nm and collecting emission (λ_em)_ at 450 and 525 nm for NAD(P)H and FAD^+^, respectively. Calibration was performed inducing maximal oxidation and reduction of NAD(P)H/FADH2 with FCCP (5 µmol/L) and cyanide (4 mmol/L), respectively.

### Intracellular Calcium measurement

LV cardiac myocytes were isolated by enzymatic digestion and were paced by electrical field stimulation at 37°C using a customized IonOptix system as described previously(Kohlhaas and Maack, 2010; Kohlhaas et al., 2010). [Ca^2+^]_C_ was measured by incubating cells with indo-1 AM (5 μmol/L) for 20 min at 25°C (λexc=340 nm, λem=405/485 nm). The bath solution contains (mM): 130 NaCL; 5 KCl; 1 MgCL2; 10 sodium HEPES; 2 sodium pyruvate; 0.3 ascorbic acid; 10 glucose and was adjust at pH 7.4 at 37°C; Isoproterenol was used at 3×10^-8^ M. The Ca^2+^ transients were analysed with the IONWIZRD software from IONOptix. Data were collected and statistically analysed using paired or unpaired tTest or 2way ANOVA & Bonferronís Multiple Comparison Test.

### Mitochondrial swelling and mPTP opening

Calcium-induced mitochondrial permeability transition pore (mPTP) opening was determined in freshly isolated cardiac mitochondria from WT and cMKO mice aged 8-10 weeks as previously described (Palmeira CM, 2018). Mitochondria were suspended in Ca^2+^ uptake buffer pH 7.4 (120 mM KCl, 5 mM MOPS, 5 mM KH_2_PO_4_, 10 mM Glutamate, 5 mM Malate *plus* cOmplete™, EDTA-free Protease Inhibitor Cocktail (Roche) at a concentration of 0.5 mg/mL and stimulated by the addition of a single pulse of 120 μM CaCl_2_.

mPTP opening was assessed in mitochondria of WT, *Mtfp1*^-/-^, *Ppif*^-/-^ and *Mtfp1*^-/-^*Ppif*^-/-^ MEFs as previously described (Karch et al., 2013) at a concentration of 1 mg/mL in mitochondrial swelling buffer (120 mM KCl, 10 mM Tris pH 7.4, 5 mM KH_2_PO_4_, 7 mM pyruvate, 1 mM malate, and 10 μM EDTA). Swelling was induced by pulsing mitochondria with 250 μM CaCl_2_ in EDTA-free buffer.

The absorbance (540 nm) was measured at intervals of ∼20 seconds at 37°C using the microplate reader Infinite® M Plex (Tecan). Cyclosporin A (CsA, 1 μM Sigma) was used as a control to inhibit mPTP-dependent mitochondrial swelling.

### BN-PAGE of cardiac mitochondria

BN-PAGE was performed as described previously (Wittig et al., 2006) with some modifications. Briefly, cardiac mitochondria were isolated from WT and cMKO mice at 8-10 weeks and solubilized with sample buffer (30 mM HEPES, 150 mM Potassium Acetate, 12% glycerol, 2 mM acid 6-aminohexanoic, 1 mM disodium EDTA) containing digitonin 6% (w/w). Lysates were incubated for 1h at 4°C and then centrifuged at 30,000 g for 20 minutes. Supernatants were then mixed with G250 sample additive 1% (Invitrogen, BN2004) and resolved on 3-12% Bis-Tris Gels (1.0 mm) (Invitrogen, BN2011BX10) using the anode running buffer (Invitrogen, BN2001) and Cathode Buffer Additive added to the anode buffer (InVitroGen, BN2002). Gels were then subjected to a semi-dry transfer using PVDF membranes (GE, 10600023). Membranes were washed with methanol and then blocked for at least 1h with 5% (w/v) semi-skimmed dry milk dissolved in Tris-buffered saline Tween 0.1% (TBST), incubated overnight at 4°C with primary antibodies dissolved 1:1.000 in 2% (w/v) Bovine Serum Albumin (BSA), 0.1% TBST. The next day membranes were incubated in secondary antibodies conjugated to horseradish peroxidase (HRP) at room temperature for 2h (diluted 1:10,000 in BSA 2% TBST 0.1%). Finally, membranes were incubated in Clarity™ Western ECL Substrate (Bio-Rad) for 2 min and luminescence was detected using the ChemiDoc® Gel Imaging System. Primary antibodies are listed in Dataset EV4.

### Co-Immunoprecipitation assay

500 µg of cardiac crude mitochondria were freshly isolated from heart tissue of cardiomyocyte specific Flag-MTFP1 Knock-In (KI) mice (*Myh6-Cre^tg/+^Mtfp1^+/+^, CAG^tg+/^)* and WT mice (*Myh6-Cre^+/+^Mtfp1^+/+^, CAG^tg/+^)* as described above. Mitochondria were lysed in IP buffer (20 mM HEPES-KOH pH 7.5, 150 mM NaCl, 0.25% Triton X-100, protease inhibitor cocktail) on ice for 20 min and then centrifugated at 10000 g, 4°C for 15 min. Supernatant obtained by centrifugation was then incubated with 20 μL of anti-FLAG magnetic beads (Sigma M8823) for 2 hours at 4°C. The immunocomplexes were then washed with IP buffer without Triton X-100 and eluted with Laemmli Sample Buffer 2x at 95°C for 5 min. Protein were stacked in a 15 % SDS-PAGE gel with a 10 min long migration at 80 V. Proteins were fixed in gel and migration was visualized using the Instant Blue stain (Expedeon). Bands were excised for digestion. Gel bands were washed twice in Ammonium bicarbonate (AmBi) 50 mM, once with AmBi 50 mM / ACN 50 % and once with 100% ANC. Gel band were incubated for 30 min at 56°C in 5 mM dithiothreitol (DTT) solution for reduction. Gel bands were washed in AmBi 50 mM and then in 100% ACN. Alkylation was performed at room temp in the dark by incubation of the gel bands in Iodocateamide 55 mM solution. Gel bands were washed twice in AmBi 50mM and in 100% ACN. 600 ng of trypsin were added for a 8h digestion at 37°C. Peptides were extracted by collecting 3 washes of the gel bands using AmBi 50 mM / 50 % ACN and 5 % FA. Peptides clean up and desalting was done using Stage tips (2 disc Empore C18 discs stacked in a P200 tip).

LC-MS/SM analysis of digested peptides was performed on an Orbitrap Q Exactive HF mass spectrometer (Thermo Fisher Scientific, Bremen) coupled to an EASY-nLC 1200 (Thermo Fisher Scientific). A home-made column was used for peptide separation (C_18_ 30 cm capillary column picotip silica emitter tip (75 μm diameter filled with 1.9 μm Reprosil-Pur Basic C_18_-HD resin, (Dr. Maisch GmbH, Ammerbuch-Entringen, Germany)). It was equilibrated and peptide were loaded in solvent A (0.1 % FA) at 900 bars. Peptides were separated at 250 nl.min^-1^. Peptides were eluted using a gradient of solvent B (ACN, 0.1 % FA) from 3% to 26% in 105 min, 26% to 48% in 20 min (total length of the chromatographic run was 145 min including high ACN level step and column regeneration). Mass spectra were acquired in data-dependent acquisition mode with the XCalibur 2.2 software (Thermo Fisher Scientific, Bremen) with automatic switching between MS and MS/MS scans using a top 12 method. MS spectra were acquired at a resolution of 60000 (at *m/z* 400) with a target value of 3 × 10^6^ ions. The scan range was limited from 400 to 1700 *m/z*. Peptide fragmentation was performed using HCD with the energy set at 26 NCE. Intensity threshold for ions selection was set at 1 × 10^5^ ions with charge exclusion of z = 1 and z > 7. The MS/MS spectra were acquired at a resolution of 15000 (at *m/z* 400). Isolation window was set at 1.6 Th. Dynamic exclusion was employed within 30 s.

#### Data Processing

Data were searched using MaxQuant (version 1.6.6.0) [1,2] using the Andromeda search engine [3] against a reference proteome of Mus musculus (53449 entries, downloaded from Uniprot the 24^th of^ July 2018). A modified sequence of the protein MTP18 with a Flag tag in its N-ter part was also searched.

The following search parameters were applied: carbamidomethylation of cysteines was set as a fixed modification, oxidation of methionine and protein N-terminal acetylation were set as variable modifications. The mass tolerances in MS and MS/MS were set to 5 ppm and 20 ppm respectively. Maximum peptide charge was set to 7 and 5 amino acids were required as minimum peptide length. A false discovery rate of 1% was set up for both protein and peptide levels. The iBAQ intensity was used to estimate the protein abundance within a sample. The match between runs features was allowed for biological replicate only.

For this Affinity Purification Mass Spectrometry analysis part, the mass spectrometry data have been deposited at the ProteomeXchange Consortium (http://www.proteomexchange.org) via the PRIDE partner repository(Perez-Riverol et al., 2019; Vizcaíno et al., 2014) with the dataset identifier PXD028529

#### Data analysis

Quantitative analysis was based on pairwise comparison of intensities. Values were log-transformed (log2). Reverse hits and potential contaminant were removed from the analysis. Proteins with at least 2 peptides (including one unique peptide) were kept for further statistics. Intensities values were normalized by median centering within conditions (normalizeD function of the R package DAPAR). Remaining proteins without any iBAQ value in one of both conditions have been considered as proteins quantitatively present in a condition and absent in the other. They have therefore been set aside and considered as differentially abundant proteins. Next, missing values were imputed using the impute.MLE function of the R package <u>imp4p</u>. Statistical testing was conducted using a limma t-test thanks to the R package limma(Pounds and Cheng, 2006). An adaptive Benjamini-Hochberg procedure was applied on the resulting p-values thanks to the function adjust.p of R package cp4p(Smyth, 2004) using the robust method previously described(Giai Gianetto et al., 2016) to estimate the proportion of true null hypotheses among the set of statistical tests. The proteins associated to an adjusted p-value inferior to a FDR level of 1% have been considered as significantly differentially abundant proteins.

### Statistical Analyses

Experiments were repeated at least three times and quantitative analyses were conducted blindly. Randomization of groups (e.g., different genotypes) was performed when simultaneous, parallel measurements were not performed (e.g., Oroboros, CM isolation). For high-throughput measurements (e.g., mitochondrial morphology, cell death), all groups were measured in parallel to reduce experimental bias. Statistical analyses were performed using GraphPad Prism 9 software. Data are presented as mean ± SD or SEM where indicated. The statistical tests used, and value of experiment replicates are described in the figure legends. Comparisons between two groups were performed by unpaired two-tailed T test. To compare more than two groups or groups at multiple time points 1-way ANOVA or 2-way ANOVA was applied. Tests were considered significant at p-value < 0.05 (*p< 0.05; **p< 0.01; ***p < 0.001; ****p< 0.0001).

## Acknowledgements

We thank Pierre-Henri Commere and Sandrine Schmutz for flow cytometry services at the Institut Pasteur and Anastasia Gazi for electron microscopy services. We thank Corinne Lesaffre at the Paris Cardiovascular Research Center for the histology services and Anu Susan Kurian and Priscilla Lopes for technical assistance. We thank Sylvie Fabrega of the Viral Vector for Gene Transfer core facility of Structure Fédérative de Recherche Necker, Université de Paris for lentiviral particle synthesis and Nils-Göran Larsson for providing mitoYFP mice. We thank Arnaud Mourier for inciteful discussions on bioenergetics and Marie Lemesle for excellent administrative assistance. T.W. is supported by the European Research Council (ERC) Starting Grant No. 714472 (Acronym “*Mitomorphosis*”), Fondation pour la Recherche Medicale (MND202003011475), and the Agence Nationale pour la Recherche (ANR-20-CE14-0039-02). C. M. is supported by the German Research Foundation (DFG; SFB 894, TRR-219; Ma 2528/7-1) and the German Federal Ministry of Education and Research (BMBF; 01EO1504).

## References

Acin-Perez, R., Lechuga-Vieco, A.V., Muñoz, M. del M., Nieto-Arellano, R., Torroja, C., Sánchez-Cabo, F., Jiménez, C., González-Guerra, A., Carrascoso, I., Benincá, C., et al. (2018). Ablation of the stress protease OMA1 protects against heart failure in mice. Science Translational Medicine 10.

Ackers-Johnson, M., Li, P.Y., and Holmes, A.P. (2016). A Simplified, Langendorff-Free Method for Concomitant Isolation of Viable Cardiac Myocytes and Non-Myocytes from the Adult Mouse Heart. Circulation.

Adams, S.H. (2000). Uncoupling protein homologs: emerging views of physiological function. J. Nutr. 130, 711–714.

Agah, R., Frenkel, P.A., French, B.A., Michael, L.H., Overbeek, P.A., and Schneider, M.D. (1997). Gene recombination in postmitotic cells. Targeted expression of Cre recombinase provokes cardiac-restricted, site-specific rearrangement in adult ventricular muscle in vivo. J. Clin. Invest. 100, 169–179.

Anand, R., Wai, T., Baker, M.J., Kladt, N., Schauss, A.C., Rugarli, E., and Langer, T. (2014). The i-AAA protease YME1L and OMA1 cleave OPA1 to balance mitochondrial fusion and fission. The Journal of Cell Biology 204, 919–929.

Antonicka, H., Leary, S.C., Guercin, G.-H., Agar, J.N., Horvath, R., Kennaway, N.G., Harding, C.O., Jaksch, M., and Shoubridge, E.A. (2003a). Mutations in COX10 result in a defect in mitochondrial heme A biosynthesis and account for multiple, early-onset clinical phenotypes associated with isolated COX deficiency. Hum. Mol. Genet. 12, 2693–2702.

Antonicka, H., Mattman, A., Carlson, C.G., Glerum, D.M., Hoffbuhr, K.C., Leary, S.C., Kennaway, N.G., and Shoubridge, E.A. (2003b). Mutations in COX15 Produce a Defect in the Mitochondrial Heme Biosynthetic Pathway, Causing Early-Onset Fatal Hypertrophic Cardiomyopathy. The American Journal of Human Genetics 72, 101–114.

Ashrafian, H., Docherty, L., Leo, V., Towlson, C., Neilan, M., Steeples, V., Lygate, C.A., Hough, T., Townsend, S., Williams, D., et al. (2010). A Mutation in the Mitochondrial Fission Gene Dnm1l Leads to Cardiomyopathy. PLOS Genetics 6, e1001000.

Aung, L.H.H., Li, R., Prabhakar, B.S., and Li, P. (2017a). Knockdown of Mtfp1 can minimize doxorubicin cardiotoxicity by inhibiting Dnm1l-mediated mitochondrial fission. Journal of Cellular and Molecular Medicine 21, 3394–3404.

Aung, L.H.H., Li, R., Prabhakar, B.S., Maker, A.V., and Li, P. (2017b). Mitochondrial protein 18 (MTP18) plays a pro-apoptotic role in chemotherapy-induced gastric cancer cell apoptosis. Oncotarget.

Aung, L.H.H., Li, Y.-Z., Yu, H., Chen, X., Yu, Z., Gao, J., and Li, P. (2019). Mitochondrial protein 18 is a positive apoptotic regulator in cardiomyocytes under oxidative stress. Clin. Sci. 133, 1067–1084.

Baines, C.P., Kaiser, R.A., Purcell, N.H., Blair, N.S., Osinska, H., Hambleton, M.A., Brunskill, E.W., Sayen, M.R., Gottlieb, R.A., Dorn, G.W., et al. (2005). Loss of cyclophilin D reveals a critical role for mitochondrial permeability transition in cell death. Nature 434, 658–662.

Basso, E., Petronilli, V., Forte, M.A., and Bernardi, P. (2008). Phosphate is essential for inhibition of the mitochondrial permeability transition pore by cyclosporin A and by cyclophilin D ablation. J. Biol. Chem. 283, 26307–26311.

Bernardi, P., Rasola, A., Forte, M., and Lippe, G. (2015). The Mitochondrial Permeability Transition Pore: Channel Formation by F-ATP Synthase, Integration in Signal Transduction, and Role in Pathophysiology. Physiol. Rev. 95, 1111–1155.

Bertero, E., Nickel, A., Kohlhaas, M., Hohl, M., Sequeira, V., Brune, C., Schwemmlein, J., Abeßer, M., Schuh, K., Kutschka, I., et al. Loss of Mitochondrial Ca2+ Uniporter Limits Inotropic Reserve and Provides Trigger and Substrate for Arrhythmias in Barth Syndrome Cardiomyopathy. Circulation 0.

Bertholet, A.M., Chouchani, E.T., Kazak, L., Angelin, A., Fedorenko, A., Long, J.Z., Vidoni, S., Garrity, R., Cho, J., Terada, N., et al. (2019). H+ transport is an integral function of the mitochondrial ADP/ATP carrier. Nature 571, 515–520.

Bock, F.J., and Tait, S.W.G. (2020). Mitochondria as multifaceted regulators of cell death. Nature Reviews Molecular Cell Biology 21, 85–100.

Brand, M.D., Affourtit, C., Esteves, T.C., Green, K., Lambert, A.J., Miwa, S., Pakay, J.L., and Parker, N. (2004). Mitochondrial superoxide: production, biological effects, and activation of uncoupling proteins. Free Radic. Biol. Med. 37, 755–767.

Brand, M.D., Pakay, J.L., Ocloo, A., Kokoszka, J., Wallace, D.C., Brookes, P.S., and Cornwall, E.J. (2005). The basal proton conductance of mitochondria depends on adenine nucleotide translocase content. Biochemical Journal 392, 353–362.

Burke, M.A., Chang, S., Wakimoto, H., Gorham, J.M., Conner, D.A., Christodoulou, D.C., Parfenov, M.G., DePalma, S.R., Eminaga, S., Konno, T., et al. (2016). Molecular profiling of dilated cardiomyopathy that progresses to heart failure. JCI Insight 1.

Carraro, M., Checchetto, V., Sartori, G., Kucharczyk, R., di Rago, J.-P., Minervini, G., Franchin, C., Arrigoni, G., Giorgio, V., Petronilli, V., et al. (2018). High-Conductance Channel Formation in Yeast Mitochondria is Mediated by F-ATP Synthase e and g Subunits. Cell. Physiol. Biochem. 50, 1840–1855.

Carraro, M., Carrer, A., Urbani, A., and Bernardi, P. (2020). Molecular nature and regulation of the mitochondrial permeability transition pore(s), drug target(s) in cardioprotection. Journal of Molecular and Cellular Cardiology 144, 76–86.

Chance, B., and Williams, G.R. (1956). The Respiratory Chain and Oxidative Phosphorylation. In Advances in Enzymology and Related Areas of Molecular Biology, (John Wiley & Sons, Ltd), pp. 65–134.

Chen, E.Y., Tan, C.M., Kou, Y., Duan, Q., Wang, Z., Meirelles, G.V., Clark, N.R., and Ma’ayan, A. (2013). Enrichr: interactive and collaborative HTML5 gene list enrichment analysis tool. BMC Bioinformatics 14, 128.

Chen, H., Ren, S., Clish, C., Jain, M., Mootha, V., McCaffery, J.M., and Chan, D.C. (2015). Titration of mitochondrial fusion rescues Mff-deficient cardiomyopathy. J. Cell Biol. 211, 795–805.

Chen, L., Gong, Q., Stice, J.P., and Knowlton, A.A. (2009). Mitochondrial OPA1, apoptosis, and heart failure. Cardiovascular Research 84, 91–99.

Chen, Y., Csordás, G., Jowdy, C., Schneider, T.G., Csordás, N., Wang, W., Liu, Y., Kohlhaas, M., Meiser, M., Bergem, S., et al. (2012). Mitofusin 2-containing mitochondrial-reticular microdomains direct rapid cardiomyocyte bioenergetic responses via interorganelle Ca(2+) crosstalk. Circ. Res. 111, 863–875.

Chiao, Y.A., Zhang, H., Sweetwyne, M., Whitson, J., Ting, Y.S., Basisty, N., Pino, L.K., Quarles, E., Nguyen, N.-H., Campbell, M.D., et al. (2020). Late-life restoration of mitochondrial function reverses cardiac dysfunction in old mice. ELife 9, e55513.

Cho, B., Cho, H.M., Jo, Y., Kim, H.D., Song, M., Moon, C., Kim, H., Kim, K., Sesaki, H., Rhyu, I.J., et al. (2017). Constriction of the mitochondrial inner compartment is a priming event for mitochondrial division. Nature Communications 8, 15754.

Christidi, E., and Brunham, L.R. (2021). Regulated cell death pathways in doxorubicin-induced cardiotoxicity. Cell Death Dis 12, 1–15.

Cokelaer, T., Desvillechabrol, D., Legendre, R., and Cardon, M. (2017). “Sequana”: a Set of Snakemake NGS pipelines. JOSS 2, 352.

Cortassa, S., Aon, M.A., Marbán, E., Winslow, R.L., and O’Rourke, B. (2003). An Integrated Model of Cardiac Mitochondrial Energy Metabolism and Calcium Dynamics. Biophysical Journal 84, 2734–2755.

Cretin, E., Lopes, P., Vimont, E., Tatsuta, T., Langer, T., Gazi, A., Sachse, M., Yu-Wai-Man, P., Reynier, P., and Wai, T. (2021). High-throughput screening identifies suppressors of mitochondrial fragmentation in OPA1 fibroblasts. EMBO Molecular Medicine *n/a*, e13579.

Dobin, A., Davis, C.A., Schlesinger, F., Drenkow, J., Zaleski, C., Jha, S., Batut, P., Chaisson, M., and Gingeras, T.R. (2013). STAR: ultrafast universal RNA-seq aligner. Bioinformatics 29, 15–21.

Duroux-Richard, I., Roubert, C., Ammari, M., Présumey, J., Grün, J.R., Häupl, T., Grützkau, A., Lecellier, C.-H., Boitez, V., Codogno, P., et al. (2016). miR-125b controls monocyte adaptation to inflammation through mitochondrial metabolism and dynamics. Blood 128, 3125–3136.

Echtay, K.S., Roussel, D., St-Pierre, J., Jekabsons, M.B., Cadenas, S., Stuart, J.A., Harper, J.A., Roebuck, S.J., Morrison, A., Pickering, S., et al. (2002). Superoxide activates mitochondrial uncoupling proteins. Nature 415, 96–99.

Ewels, P., Magnusson, M., Lundin, S., and Käller, M. (2016). MultiQC: summarize analysis results for multiple tools and samples in a single report. Bioinformatics 32, 3047–3048.

Fernandez-Caggiano, M., Kamynina, A., Francois, A.A., Prysyazhna, O., Eykyn, T.R., Krasemann, S., Crespo-Leiro, M.G., Vieites, M.G., Bianchi, K., Morales, V., et al. (2020). Mitochondrial pyruvate carrier abundance mediates pathological cardiac hypertrophy. Nat Metab 2, 1223–1231.

Fonseca, T.B., Sánchez-Guerrero, Á., Milosevic, I., and Raimundo, N. (2019). Mitochondrial fission requires DRP1 but not dynamins. Nature 570, E34–E42.

Ghazal, N., Peoples, J.N., Mohiuddin, T.A., and Kwong, J.Q. (2021). Mitochondrial functional resilience after TFAM ablation in the adult heart. Am. J. Physiol. Cell Physiol.

Giacomello, M., Pyakurel, A., Glytsou, C., and Scorrano, L. (2020). The cell biology of mitochondrial membrane dynamics. Nature Reviews Molecular Cell Biology.

Giai Gianetto, Q., Combes, F., Ramus, C., Bruley, C., Couté, Y., and Burger, T. (2016). Calibration plot for proteomics: A graphical tool to visually check the assumptions underlying FDR control in quantitative experiments. PROTEOMICS 16, 29–32.

Graham, B.H., Waymire, K.G., Cottrell, B., Trounce, I.A., MacGregor, G.R., and Wallace, D.C. (1997). A mouse model for mitochondrial myopathy and cardiomyopathy resulting from a deficiency in the heart/muscle isoform of the adenine nucleotide translocator. Nat Genet 16, 226–234.

Halestrap, A.P., and Davidson, A.M. (1990). Inhibition of Ca2+-induced large-amplitude swelling of liver and heart mitochondria by cyclosporin is probably caused by the inhibitor binding to mitochondrial-matrix peptidyl-prolyl cis-trans isomerase and preventing it interacting with the adenine nucleotide translocase. Biochemical Journal 268, 153–160.

Hansson, A., Hance, N., Dufour, E., Rantanen, A., Hultenby, K., Clayton, D.A., Wibom, R., and Larsson, N.-G. (2004). A switch in metabolism precedes increased mitochondrial biogenesis in respiratory chain-deficient mouse hearts. Proc. Natl. Acad. Sci. U. S. A. 101, 3136–3141.

Heide, H., Bleier, L., Steger, M., Ackermann, J., Dröse, S., Schwamb, B., Zörnig, M., Reichert, A.S., Koch, I., Wittig, I., et al. (2012). Complexome profiling identifies TMEM126B as a component of the mitochondrial complex I assembly complex. Cell Metab. 16, 538–549.

Hohl, M., Wagner, M., Reil, J.-C., Müller, S.-A., Tauchnitz, M., Zimmer, A.M., Lehmann, L.H., Thiel, G., Böhm, M., Backs, J., et al. (2013). HDAC4 controls histone methylation in response to elevated cardiac load. J Clin Invest 123, 1359–1370.

Houweling, A.C., van Borren, M.M., Moorman, A.F.M., and Christoffels, V.M. (2005). Expression and regulation of the atrial natriuretic factor encoding gene Nppa during development and disease. Cardiovasc Res 67, 583–593.

Kageyama, Y., Hoshijima, M., Seo, K., Bedja, D., Sysa-Shah, P., Andrabi, S.A., Chen, W., Höke, A., Dawson, V.L., Dawson, T.M., et al. (2014). Parkin-independent mitophagy requires Drp1 and maintains the integrity of mammalian heart and brain. The EMBO Journal 33, 2798–2813.

Kang, B.H., Plescia, J., Dohi, T., Rosa, J., Doxsey, S.J., and Altieri, D.C. (2007). Regulation of Tumor Cell Mitochondrial Homeostasis by an Organelle-Specific Hsp90 Chaperone Network. Cell 131, 257–270.

Karamanlidis, G., Lee, C.F., Garcia-Menendez, L., Kolwicz, S.C., Suthammarak, W., Gong, G., Sedensky, M.M., Morgan, P.G., Wang, W., and Tian, R. (2013). Mitochondrial Complex I Deficiency Increases Protein Acetylation and Accelerates Heart Failure. Cell Metabolism 18, 239–250.

Karch, J., Kwong, J.Q., Burr, A.R., Sargent, M.A., Elrod, J.W., Peixoto, P.M., Martinez-Caballero, S., Osinska, H., Cheng, E.H.-Y., Robbins, J., et al. (2013). Bax and Bak function as the outer membrane component of the mitochondrial permeability pore in regulating necrotic cell death in mice. Elife 2, e00772.

Karch, J., Bround, M.J., Khalil, H., Sargent, M.A., Latchman, N., Terada, N., Peixoto, P.M., and Molkentin, J.D. (2019). Inhibition of mitochondrial permeability transition by deletion of the ANT family and CypD. Sci Adv 5, eaaw4597.

Kohlhaas, M., and Maack, C. (2010). Adverse Bioenergetic Consequences of Na+-Ca2+ Exchanger–Mediated Ca2+ Influx in Cardiac Myocytes. Circulation 122, 2273–2280.

Kohlhaas, M., Liu, T., Knopp, A., Zeller, T., Ong, M.F., Böhm, M., O’Rourke, B., and Maack, C. (2010). Elevated Cytosolic Na+ Increases Mitochondrial Formation of Reactive Oxygen Species in Failing Cardiac Myocytes. Circulation 121, 1606–1613.

Kohlhaas, M., Nickel, A.G., and Maack, C. (2017). Mitochondrial energetics and calcium coupling in the heart. The Journal of Physiology 595, 3753–3763.

Kolwicz, S.C., Jr, Purohit, S., and Tian, R. (2013). Cardiac metabolism and its interactions with contraction, growth, and survival of cardiomyocytes. Circ. Res. 113, 603–616.

Köster, J., and Rahmann, S. (2012). Snakemake—a scalable bioinformatics workflow engine. Bioinformatics 28, 2520–2522.

Kulak, N.A., Pichler, G., Paron, I., Nagaraj, N., and Mann, M. (2014). Minimal, encapsulated proteomic-sample processing applied to copy-number estimation in eukaryotic cells. Nat Methods 11, 319–324.

Kwong, J.Q., and Molkentin, J.D. (2015). Physiological and Pathological Roles of the Mitochondrial Permeability Transition Pore in the Heart. Cell Metabolism 21, 206–214.

Kwong, J.Q., Davis, J., Baines, C.P., Sargent, M.A., Karch, J., Wang, X., Huang, T., and Molkentin, J.D. (2014). Genetic deletion of the mitochondrial phosphate carrier desensitizes the mitochondrial permeability transition pore and causes cardiomyopathy. Cell Death Differ. 21, 1209–1217.

Lei, Y., Guerra Martinez, C., Torres-Odio, S., Bell, S.L., Birdwell, C.E., Bryant, J.D., Tong, C.W., Watson, R.O., West, L.C., and West, A.P. (2021). Elevated type I interferon responses potentiate metabolic dysfunction, inflammation, and accelerated aging in mtDNA mutator mice. Sci. Adv. 7, eabe7548.

Lewis, S.C., Uchiyama, L.F., and Nunnari, J. (2016). ER-mitochondria contacts couple mtDNA synthesis with mitochondrial division in human cells. Science 353, aaf5549–aaf5549.

Liao, X., Zhang, R., Lu, Y., Prosdocimo, D.A., Sangwung, P., Zhang, L., Zhou, G., Anand, P., Lai, L., Leone, T.C., et al. (2015). Kruppel-like factor 4 is critical for transcriptional control of cardiac mitochondrial homeostasis. J. Clin. Invest. 125, 3461–3476.

Liao, Y., Smyth, G.K., and Shi, W. (2014). featureCounts: an efficient general purpose program for assigning sequence reads to genomic features. Bioinformatics 30, 923–930.

Livak, K.J., and Schmittgen, T.D. (2001). Analysis of Relative Gene Expression Data Using Real-Time Quantitative PCR and the 2−ΔΔCT Method. Methods 25, 402–408.

Losón, O.C., Song, Z., Chen, H., and Chan, D.C. (2013). Fis1, Mff, MiD49, and MiD51 mediate Drp1 recruitment in mitochondrial fission. Mol. Biol. Cell 24, 659–667.

Lu, J.-H., Shi, Z.-F., and Xu, H. (2014). The mitochondrial cyclophilin D/p53 complexation mediates doxorubicin-induced non-apoptotic death of A549 lung cancer cells. Mol Cell Biochem 389, 17–24.

Macher, G., Koehler, M., Rupprecht, A., Kreiter, J., Hinterdorfer, P., and Pohl, E.E. (2018). Inhibition of mitochondrial UCP1 and UCP3 by purine nucleotides and phosphate. Biochim. Biophys. Acta Biomembr. 1860, 664–672.

Martin, M. (2011). Cutadapt removes adapter sequences from high-throughput sequencing reads. EMBnet.Journal 17, 10–12.

Mastronarde, D.N., and Held, S.R. (2017). Automated tilt series alignment and tomographic reconstruction in IMOD. Journal of Structural Biology 197, 102–113.

Milduberger, N., Bustos, P.L., González, C., Perrone, A.E., Postan, M., and Bua, J. (2021). Trypanosoma cruzi infection in Cyclophilin D deficient mice. Exp. Parasitol. 220, 108044.

Mitchell, P. (1961). Coupling of Phosphorylation to Electron and Hydrogen Transfer by a Chemi-Osmotic type of Mechanism. Nature 191, 144–148.

Mootha, V.K., Arai, A.E., and Balaban, R.S. (1997). Maximum oxidative phosphorylation capacity of the mammalian heart. American Journal of Physiology-Heart and Circulatory Physiology 272, H769–H775.

Morciano, G., Naumova, N., Koprowski, P., Valente, S., Sardão, V.A., Potes, Y., Rimessi, A., Wieckowski, M.R., and Oliveira, P.J. The mitochondrial permeability transition pore: an evolving concept critical for cell life and death. Biological Reviews *n/a*.

Morita, S.-Y., and Terada, T. (2015). Enzymatic measurement of phosphatidylglycerol and cardiolipin in cultured cells and mitochondria. Sci. Rep. 5, 11737.

Morita, M., Prudent, J., Basu, K., Goyon, V., Katsumura, S., Hulea, L., Pearl, D., Siddiqui, N., Strack, S., McGuirk, S., et al. (2017). mTOR Controls Mitochondrial Dynamics and Cell Survival via MTFP1. Mol Cell 67, 922–935.e5.

Mourier, A., Ruzzenente, B., Brandt, T., Kuhlbrandt, W., and Larsson, N.-G. (2014). Loss of LRPPRC causes ATP synthase deficiency. Human Molecular Genetics 23, 2580–2592.

Nakagawa, T., Shimizu, S., Watanabe, T., Yamaguchi, O., Otsu, K., Yamagata, H., Inohara, H., Kubo, T., and Tsujimoto, Y. (2005). Cyclophilin D-dependent mitochondrial permeability transition regulates some necrotic but not apoptotic cell death. Nature 434, 652–658.

Ng, M.Y.W., Wai, T., and Simonsen, A. (2021). Quality control of the mitochondrion. Developmental Cell.

Nickel, A.G., von Hardenberg, A., Hohl, M., Löffler, J.R., Kohlhaas, M., Becker, J., Reil, J.-C., Kazakov, A., Bonnekoh, J., Stadelmaier, M., et al. (2015). Reversal of Mitochondrial Transhydrogenase Causes Oxidative Stress in Heart Failure. Cell Metabolism 22, 472–484.

Osellame, L.D., Singh, A.P., Stroud, D.A., Palmer, C.S., Stojanovski, D., Ramachandran, R., and Ryan, M.T. (2016). Cooperative and independent roles of the Drp1 adaptors Mff, MiD49 and MiD51 in mitochondrial fission. J. Cell Sci. 129, 2170–2181.

Perez-Riverol, Y., Csordas, A., Bai, J., Bernal-Llinares, M., Hewapathirana, S., Kundu, D.J., Inuganti, A., Griss, J., Mayer, G., Eisenacher, M., et al. (2019). The PRIDE database and related tools and resources in 2019: improving support for quantification data. Nucleic Acids Research 47, D442–D450.

Pounds, S., and Cheng, C. (2006). Robust estimation of the false discovery rate. Bioinformatics 22, 1979–1987.

Rappsilber, J., Mann, M., and Ishihama, Y. (2007). Protocol for micro-purification, enrichment, pre-fractionation and storage of peptides for proteomics using StageTips. Nat Protoc 2, 1896–1906.

Rath, S., Sharma, R., Gupta, R., Ast, T., Chan, C., Durham, T.J., Goodman, R.P., Grabarek, Z., Haas, M.E., Hung, W.H.W., et al. (2021). MitoCarta3.0: an updated mitochondrial proteome now with sub-organelle localization and pathway annotations. Nucleic Acids Res. 49, D1541–D1547.

Reimand, J., Isserlin, R., Voisin, V., Kucera, M., Tannus-Lopes, C., Rostamianfar, A., Wadi, L., Meyer, M., Wong, J., Xu, C., et al. (2019). Pathway enrichment analysis and visualization of omics data using g:Profiler, GSEA, Cytoscape and EnrichmentMap. Nat Protoc 14, 482–517.

Roesler, A., and Kazak, L. (2020). UCP1-independent thermogenesis. Biochemical Journal 477, 709–725.

Ruprecht, J.J., and Kunji, E.R.S. (2021). Structural Mechanism of Transport of Mitochondrial Carriers. Annu. Rev. Biochem. 90, 535–558.

Schorb, M., Haberbosch, I., Hagen, W.J.H., Schwab, Y., and Mastronarde, D.N. (2019). Software tools for automated transmission electron microscopy. Nature Methods 16, 471–477.

Schwanhäusser, B., Busse, D., Li, N., Dittmar, G., Schuchhardt, J., Wolf, J., Chen, W., and Selbach, M. (2011). Global quantification of mammalian gene expression control. Nature 473, 337–342.

Schwenk, F., Baron, U., and Rajewsky, K. (1995). A cre-transgenic mouse strain for the ubiquitous deletion of loxP-flanked gene segments including deletion in germ cells. Nucleic Acids Research 23, 5080–5081.

Shirakabe, A., Zhai, P., Ikeda, Y., Saito, T., Maejima, Y., Hsu, C.-P., Nomura, M., Egashira, K., Levine, B., and Sadoshima, J. (2016). Drp1-Dependent Mitochondrial Autophagy Plays a Protective Role Against Pressure Overload-Induced Mitochondrial Dysfunction and Heart Failure. Circulation 133, 1249–1263.

Smyth, G.K. (2004). Linear Models and Empirical Bayes Methods for Assessing Differential Expression in Microarray Experiments. Statistical Applications in Genetics and Molecular Biology 3.

Song, M., Mihara, K., Chen, Y., Scorrano, L., and Dorn II, G.W. (2015). Mitochondrial Fission and Fusion Factors Reciprocally Orchestrate Mitophagic Culling in Mouse Hearts and Cultured Fibroblasts. Cell Metabolism.

Sprenger, H.-G., and Langer, T. (2019). The Good and the Bad of Mitochondrial Breakups. Trends in Cell Biology 29, 888–900.

Todisco, S., Di Noia, M.A., Onofrio, A., Parisi, G., Punzi, G., Redavid, G., De Grassi, A., and Pierri, C.L. (2016). Identification of new highly selective inhibitors of the human ADP/ATP carriers by molecular docking and in vitro transport assays. Biochem. Pharmacol. 100, 112–132.

Tondera, D. (2005). The mitochondrial protein MTP18 contributes to mitochondrial fission in mammalian cells. J Cell Sci 118, 3049–3059.

Tondera, D., Santel, A., Schwarzer, R., Dames, S., Giese, K., Klippel, A., and Kaufmann, J. (2004). Knockdown of MTP18, a novel phosphatidylinositol 3-kinase-dependent protein, affects mitochondrial morphology and induces apoptosis. J. Biol. Chem. 279, 31544–31555.

Tondera, D., Grandemange, S., Jourdain, A., Karbowski, M., Mattenberger, Y., Herzig, S., Da Cruz, S., Clerc, P., Raschke, I., Merkwirth, C., et al. (2009). SLP-2 is required for stress-induced mitochondrial hyperfusion. EMBO J. 28, 1589–1600.

Tyanova, S., Temu, T., and Cox, J. (2016). The MaxQuant computational platform for mass spectrometry-based shotgun proteomics. Nat Protoc 11, 2301–2319.

Varet, H., Brillet-Guéguen, L., Coppée, J.-Y., and Dillies, M.-A. (2016). SARTools: A DESeq2-and EdgeR-Based R Pipeline for Comprehensive Differential Analysis of RNA-Seq Data. PLOS ONE 11, e0157022.

Vaseva, A.V., Marchenko, N.D., Ji, K., Tsirka, S.E., Holzmann, S., and Moll, U.M. (2012). p53 Opens the Mitochondrial Permeability Transition Pore to Trigger Necrosis. Cell 149, 1536–1548.

Vizcaíno, J.A., Deutsch, E.W., Wang, R., Csordas, A., Reisinger, F., Ríos, D., Dianes, J.A., Sun, Z., Farrah, T., Bandeira, N., et al. (2014). ProteomeXchange provides globally coordinated proteomics data submission and dissemination. Nat Biotechnol 32, 223–226.

Wai, T., and Langer, T. (2016). Mitochondrial Dynamics and Metabolic Regulation. Trends Endocrinol. Metab. 27, 105–117.

Wai, T., Garcia-Prieto, J., Baker, M.J., Merkwirth, C., Benit, P., Rustin, P., Ruperez, F.J., Barbas, C., Ibanez, B., and Langer, T. (2015). Imbalanced OPA1 processing and mitochondrial fragmentation cause heart failure in mice. Science 350, aad0116–aad0116.

Wang, K., Gan, T.-Y., Li, N., Liu, C.-Y., Zhou, L.-Y., Gao, J.-N., Chen, C., Yan, K.-W., Ponnusamy, M., Zhang, Y.-H., et al. (2017). Circular RNA mediates cardiomyocyte death via miRNA-dependent upregulation of MTP18 expression. Cell Death Differ. 24, 1111–1120.

Wittig, I., Braun, H.-P., and Schägger, H. (2006). Blue native PAGE. Nat. Protoc. 1, 418–428.

Woyda-Ploszczyca, A.M., and Jarmuszkiewicz, W. (2014). Different effects of guanine nucleotides (GDP and GTP) on protein-mediated mitochondrial proton leak. PLoS One 9, e98969.

Zhang, H., Alder, N.N., Wang, W., Szeto, H., Marcinek, D.J., and Rabinovitch, P.S. (2020a). Reduction of elevated proton leak rejuvenates mitochondria in the aged cardiomyocyte. ELife 9, e60827.

Zhang, Y., Taufalele, P.V., Cochran, J.D., Robillard-Frayne, I., Marx, J.M., Soto, J., Rauckhorst, A.J., Tayyari, F., Pewa, A.D., Gray, L.R., et al. (2020b). Mitochondrial pyruvate carriers are required for myocardial stress adaptation. Nat Metab 2, 1248–1264.

Zhou, B., and Tian, R. (2018). Mitochondrial dysfunction in pathophysiology of heart failure. J. Clin. Invest. 128, 3716–3726.

